# JIND-Multi: Leveraging Multiple Labeled Datasets for Automated Annotation of Single-Cell RNA and ATAC Data

**DOI:** 10.1101/2025.01.15.633130

**Authors:** Joseba Sancho, Akash Kanhirodan, Xabier Garrote, Olivier Gevaert, Mikel Hernaez, Guillermo Serrano, Idoia Ochoa

## Abstract

**Background:** The creation of single-cell atlases is essential for understanding cellular diversity and heterogeneity. However, assembling these atlases is challenging due to batch effects and the need for accurate cell annotation. Current methods for single-cell RNA and ATAC sequencing, while effective for integration, are not optimized for cell annotation. Additionally, many annotation tools rely on external databases or reference scRNA-Seq datasets, which may limit their adaptability to specific study needs, especially for rare cell-types or scATAC-Seq data.

**Results:** We introduce JIND-Multi, an extended version of the JIND framework, designed to transfer cell-type labels across multiple annotated datasets. JIND-Multi significantly reduces the proportion of unclassified cells in single-cell RNA sequencing (scRNA-Seq) data while maintaining the accuracy and performance of the original JIND model. Furthermore, JIND-Multi demonstrates robust and precise annotation results in its inaugural application to scATAC-Seq data, proving its versatility and effectiveness across different single-cell sequencing technologies.

**Conclusions:** JIND-Multi represents an improvement in cell annotation, reducing unassigned cells and offering a reliable solution for both scRNA-Seq and scATAC-Seq data. Its ability to handle multiple labeled datasets enhances the precision of annotations, making it a valuable tool for the single-cell research community. JIND-Multi is publicly available at:https://github.com/ML4BM-Lab/JIND-Multi.git.

## Background

The creation of single-cell atlases is crucial to modern biology and medicine, as it provides a high-resolution and comprehensive view of the diversity and heterogeneity within tissues and organisms at the individual cell level [1]. However, there is still no unified criterion for generating these atlases, and they are primarily based on single-cell RNA sequencing (scRNA-Seq) [2]. The atlas-assembling process remains time-consuming and labor-intensive, as samples from different individuals are normally sequenced over time and across different platforms, often resulting in datasets that incorporate pronounced batch effects. These effects require rigorous annotation with standardized criteria for effective integration [2].

Various integration methods have been proven effective, such as Harmony [3] and scVI [4] for scRNA-Seq, while Seurat [5] and LIGER [6] can be used for both scRNA-Seq and single-cell ATAC sequencing (scATAC-Seq) data. These methods are optimized to preserve data variability and integrate cells by removing technical noise while retaining as much original biological signal as possible. However, their integration dimension transformations are not specifically designed for the task of cell annotation [7]. In this context, several automatic annotators for scRNA-Seq have been developed, though they face several challenges. Many of these methods depend on open-access databases, like CellTypist [8], scQuery [9], and Azimuth [10], and while these databases are accessible, their preselected annotations may not address specific study needs, especially concerning annotation depth and rare cell-types. Moreover, some of these methods are designed to work exclusively with specific integration methods, which can limit their flexibility and applicability [2].

In addition, some methods use marker genes for annotation assistance, such as scQuery [9] and ScType [11], or generate an embedded space not specifically optimized for annotation, focusing instead on calculating cell-type landmarks (MARS) [12]. In contrast, the JIND framework [7] provides a more effective solution by leveraging user-specific datasets for the annotation of scRNA-Seq data. Although it is not designed to preserve all features intact, JIND is capable of integrating batches and eliminating batch effects by generating a latent space optimized for cell annotation, with label transfer performed using a unified standard criterion.

Regarding scATAC-Seq, there are fewer well-established methods. For example, there are specific tools such as AtacAnnoR [13] and SANGO [14], as well as databases like Azimuth [10]. These methods mainly depend on a scRNA-Seq reference [13] [10] or encode DNA fragments from ATAC peaks as one-hot inputs [14]. Given that real-world experiments generally provide data from one type—scRNA-Seq or scATAC-Seq—methods requiring scRNA-Seq to annotate scATAC-Seq data may have access only to unpaired scRNA-Seq data, which can affect their performance.

In light of these challenges, in this work we present JIND-Multi, an extension of the JIND framework, that allows to transfer cell-type labels from several annotated datasets, e.g., those that compose an atlas. Notably, JIND-Multi is also applicable for the annotation of scATAC-Seq data. Hence, JIND-Multi can automatically annotate scRNA-Seq data using one or more annotated scRNA-Seq batches, and scATAC-Seq data using one or more annotated scATAC-Seq batches. Similarly to its predecessor, JIND-Multi has the option to mark cells as “unassigned” if the model does not produce reliable predictions, i.e., below some pre-computed cell-type specific thresholds.

We show that by leveraging a large number of annotated datasets, JIND-Multi makes the annotation of unlabeled datasets more precise and with lower rejection rates (unassigned cells). We provide an efficient implementation of JIND-Multi that is publicly available and ready to use by the community.

## Findings

### JIND-Multi

Next, we outline the proposed method, JIND-Multi (see Supplementary Section 1 for a detailed description). The method consists of four main parts: initialization of a latent space (*h*) optimized for cell-type classification using a labeled dataset, integration of additional labeled datasets into the model’s latent space, generation of a batch-corrected latent space for the unlabeled dataset (target batch), and finally, cell-type inference by leveraging information from the multiple labeled datasets (see Figure 1).

**Figure 1.**
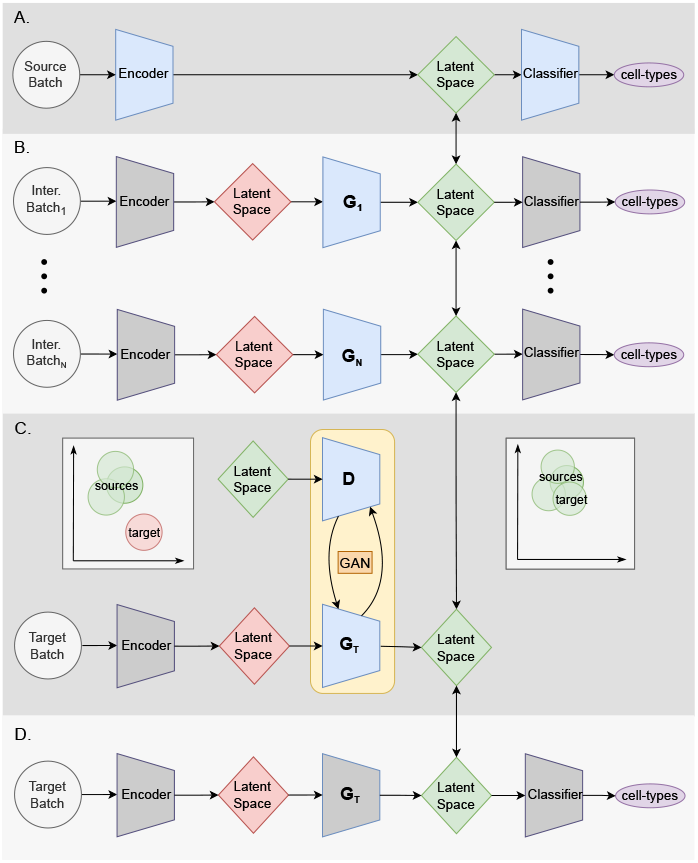
Simplified JIND-Multi workflow. **A.***Latent space initialization and classifier training*: A neural network (NN) based encoder and cell-type classifier are trained on the source dataset (colored blue). This step creates a latent space (green) suitable for cell-type classification and fixes the encoder and classifier for the remaining steps (colored grey). **B**. *Integrating additional labeled datasets into the latent space*: For each additional annotated dataset, the encoder generates a latent space with batch effect (red). To remove this noise, a NN-based generator is trained for each dataset to generate a latent space (green) that can be correctly classified by the already trained classifier, and hence that is aligned to that of the source’s. This step increases the number of samples that can be used to train the Generative Adversarial Network (GAN) of the next step. **C** *Target batch integration* and **D** *cell-type inference*: To annotate the unlabeled dataset (target), a GAN is trained such that the generator produces a latent space indistinguishable from that of the sources’ and that can be used as input to the trained classifier. More detailed figure is provided in Figure S1

JIND-Multi uses a labeled dataset as the source to initialize the model. Ideally, this source dataset should be the best representative of the data, i.e., containing all the cell-types and enough cells. If no preference is provided for the source batch selection, JIND-Multi can automatically select the source batch that minimizes the rejection rate on the target dataset (see Supplementary Section 1.1 for further details).

The initialization step involves training an encoder, which generates an embedding of the input (denoted as latent space) of reduced dimension, followed by a classifier that makes the cell-type predictions. The encoder and the classifier are based on neural networks (NN), and their architecture is as defined in Supplementary Section 1.1.

The prediction model is trained on the source batch (Figure 1A), as in JIND, by minimizing a weighted categorical cross-entropy loss (see Supplementary Section 1.1). The source batch is split into training (80%) and validation (20%) sets, using a stratified partitioning approach to ensure that the distribution of cell-types remains balanced across both subsets.

To use the additional labeled datasets (intermediate datasets) for training, JIND-Multi first aligns them to the source batch to correct potential batch effects. This alignment is performed by feeding each intermediate dataset *j* (*j* = 1, …, *n*) to the fixed (trained) encoder, which generates latent spaces *h* that include batch effects (colored red in Figure 1B). To mitigate these effects, a NN-based generator *G*_*j*_ is trained sequentially for each intermediate dataset.

The generator consists of two neural network blocks, *S*^*j*^ and *B*^*j*^, as in JIND [7], which take the gene expression vector of each cell as input and modify the latent space by adjusting the necessary scale and bias parameters. The operation is formulated as follows (see Supplementary Section 1.2 and Figure S1 for a more detailed explanation of the generator’s architecture and training):

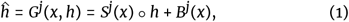

where *○* denotes element-wise multiplication.

Since the intermediate datasets are annotated, the generators are trained similarly to the previous step, minimizing the weighted cross-entropy loss, with the weights of the encoder and classifier fixed. Once the generators are trained, the resulting latent spaces *h*^ are aligned with that of the source batch (colored green in Figure 1B).

After adapting an intermediate batch to the source latent space, parallel fine-tuning is performed on the encoder and classifier using the aligned samples from different batches. During this process, the generator blocks (*S*^*j*^ and *B*^*j*^) for each intermediate dataset remain fixed, while the encoder and classifier are refined to generalize effectively across all labeled datasets. A subset comprising 10% of the intermediate dataset is used to validate the performance of this integration. This step is repeated *n* times, ensuring consistency in the latent space representation, enhancing the model’s ability to handle variability across batches, and accommodating cell types that are underrepresented in the latent space initialized by the source batch.

As part of the training process, cell-type specific thresholds are computed to avoid annotating cells with low-confidence predictions. Cell predictions below these thresholds will be marked as “unassigned” during the annotation process.

In the inference stage (Figure 1C-D), JIND-Multi aims to predict the cell-types for the unlabeled dataset, referred to as the target batch. To address batch effects in the target batch, JIND-Multi aligns its latent space with that of the annotated datasets (sources).

This alignment is achieved through the competitive training of a Generative Adversarial Network (GAN), composed of a generator *G*_*t*_ and a discriminator *D* (Figure 1C).

The discriminator *D* is a NN classifier designed to distinguish between latent spaces originating from the source batches (including the intermediate datasets) and those from the target batch. In contrast, the generator *G*_*t*_, with the same two-block structure *S*^*j*^ and *B*^*j*^ explained above, creates a modified latent code *h*^ for the target batch, aligning it more closely with the source latent space, making it indistinguishable to the discriminator. This process ensures the target batch’s latent space is effectively integrated with that of the annotated datasets.

All annotated samples are used during GAN training. In each epoch, the latent codes from the target batch are passed through the discriminator *D*, along with latent codes from the annotated batches. The parameters of the encoder and generator 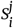 and 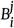 for *j* ∈ [1 : *n*] keep fixed during training. For more information about the training and architecture, refer to Supplementary Section 1.3 and Figure S1.

Once the target batch’s latent space is aligned, the trained cell-type classifier is used to predict the cell-types of the unlabeled dataset (Figure 1D).

JIND-Multi leverages information from more samples to improve the training of the GAN, enhancing its ability to align latent spaces, and ultimately increasing the accuracy and reducing the rejection rates. A final fine-tuning is performed using highconfidence predictions (see Supplementary Section 1.3 and Figure S1).

### Datasets

In this study, we employed three scRNA-Seq datasets: *Pancreas* [15], [16], [17], non-small cell lung cancer (*NSCLC Lung*) [18], and *Brain Neurips* [19], along with three scATAC-Seq datasets: human bone marrow mononuclear cells (*BMMC*) [20], *Fetal Heart*, and *Fetal Kidney* [21]. Both scRNA-Seq and scATAC-Seq data were represented as gene-by-cell matrices before processing. The data underwent count normalization, log transformation, and selection of the top 5K genes with the highest variance across batches (see Supplementary Section 2.1).

A minimum number of cells per cell-type was required (set to 100 by default), and datasets were filtered to ensure consistent cell-types across batches. Table 1 summarizes the batches, cells, and cell-types used. All considered methods were trained on normalized data, focusing on the largest batches with all cell-types present. Supplementary Section 2.2 provides further details, including download links and batch selection.

**Table 1.**
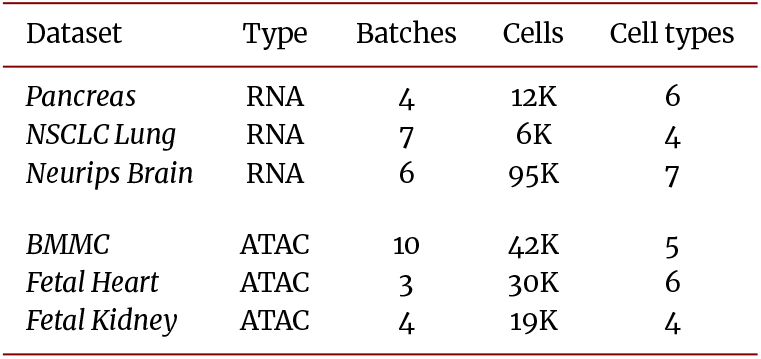
Summary of the scRNA-Seq and scATAC-Seq datasets.

To assess batch effects, UMAP [22] was used to visualize gene profiles, with cells colored by batch and cell-type (Figure S2-S7). The *Pancreas* dataset showed the most noticeable batch effect (Figure S2), while *NSCLC Lung* exhibited minimal batch variation (Figure S3).

### Experimental setup and evaluation metrics

Since JIND-Multi can train with a varying number of annotated batches, for each considered dataset we start with just the source and target batches, which is equivalent to using JIND, and then we keep adding intermediate datasets. This allows us to evaluate how the addition of labeled batches affects the prediction performance of JIND-Multi. For comparison, for each of these combinations, we also consider merging all annotated datasets into one, without correcting for batch effect, and using JIND to annotate the target batch.

In addition to JIND, we consider MARS [12] for the annotation of scRNA-Seq, as it can also leverage several annotated datasets for training, and AtacAnnoR [13] for scATAC-Seq. We note that MARS and AtacAnnoR do not incorporate a filtering scheme to reject lowconfidence predictions, and that AtacAnnoR needs a scRNA-Seq dataset in order to be able to transfer the labels based on anchor-points or gene markers. Additionally, AtacAnnoR cannot leverage more than one batch for label transfer.

Each annotation process was repeated 10 times to assess the robustness of the methods. The evaluation metrics take into account the accuracy before and after applying the filtering scheme, denoted as raw accuracy and effective accuracy, respectively, the rejection rate, i.e., the proportion of cells marked as “unassigned”, as well as the execution times.

All experiments were conducted in a workstation with 64 cores Intel xeon gold 6130 2.1GHz and 754 GB of RAM. A Quadro RTX 4000 GPU was also used, with driver version 460.67 and cuda 11.2 version.

## Results

Detailed results for the considered scRNA-Seq and scATAC-Seq datasets using a different number of intermediate batches for JIND and JIND-Multi, as well as a the comparison to MARS and AtacAnnoR is provided in Tables S7-S12. Results are summarized in Figure 2.

**Figure 2.**
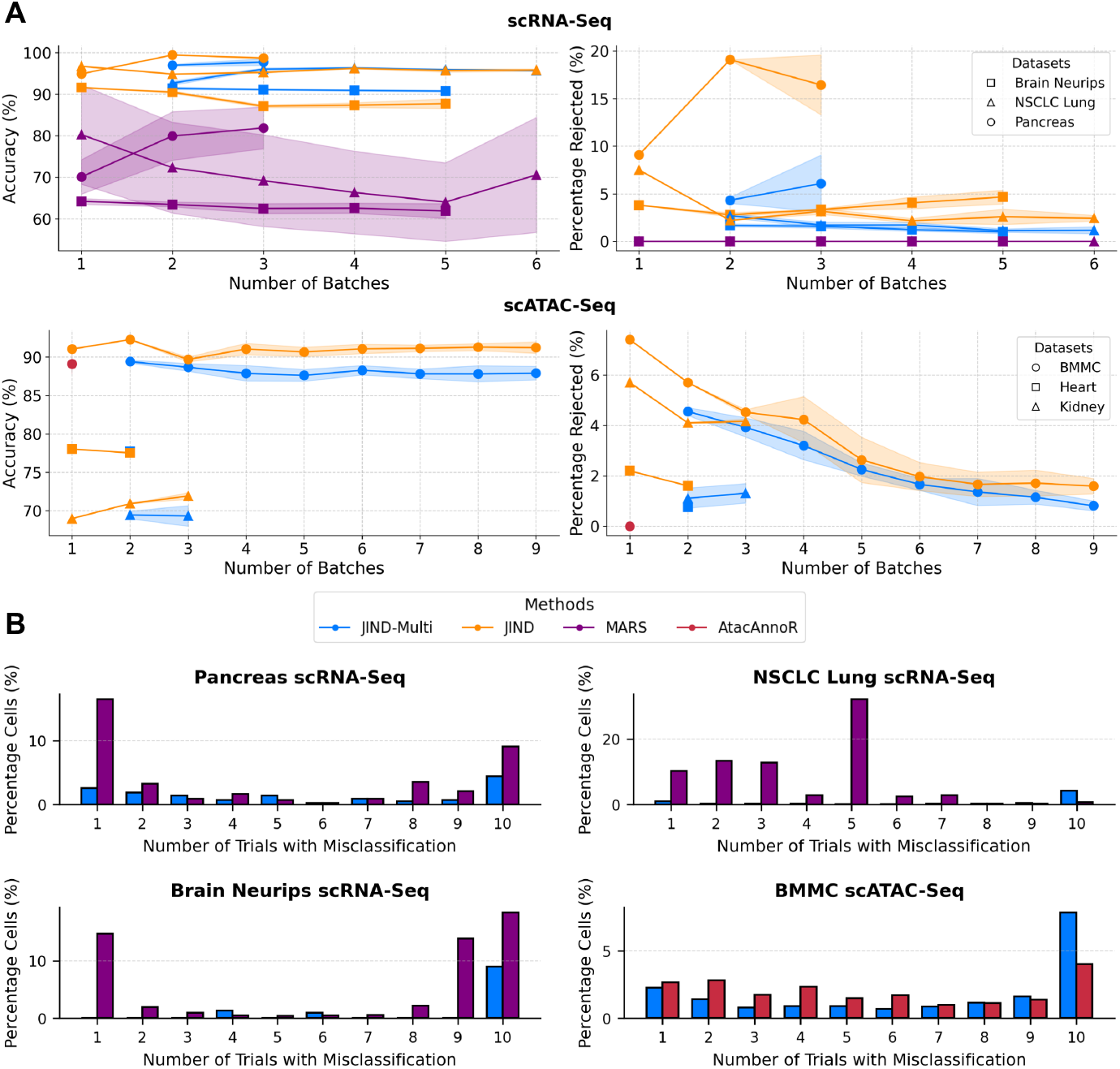
**A**. Comparative analysis of average accuracy and rejection rate across 10 iterations and employing different batch sizes on scRNA-Seq (top) and scATAC-Seq (bottom) datasets. The figure evaluates the performance of JIND (orange) and JIND-Multi (blue) across all datasets. For scRNA-Seq, we also include results for MARS (purple), and for scATAC-Seq, the comparison includes AtacAnnoR (red). For these two methods we report average raw accuracy as they do not include a filtering scheme. Datasets are marked with distinct shapes, and standard deviation is indicated by error tolerances. **B**. Bar chart comparing the percentage of cells associated with each incorrect trial count (i.e., the number of trials in which predictions were incorrect) for JIND-Multi (blue) and MARS (purple) on the thre scRNA-Seq datasets and *BMMC scATAC-Seq*. The *Pancreas scRNA-Seq* dataset uses batches *0-1-2* for both JIND-Multi and MARS, while the *Lung scRNA-Seq* dataset utilizes batches D5-D0-D1-D3-D4-D6 for both methods. For the *Brain scRNA-Seq* dataset, JIND-Multi uses batches C4-AD2, whereas MARS uses only batch C4. In the case of *BMMC*, JIND-Multi was trained with batches s4d8-s1d1-s1d2-s1d3-s2d1-s2d4-s2d5-s3d10-s4d1, while AtacAnnoR used batch s4d8.

### Annotation of scRNA-Seq data

Overall, JIND-Multi consistently matches or exceeds the performance of its predecessor JIND, achieving over 90% in both raw and effective mean accuracy while significantly reducing the rejection rate With respect to MARS, both JIND and JIND-Multi exhibit superior performance, with MARS just obtaining around 80% at most for the *Pancreas* and *NSCLC Lung* datasets, and about 65% for *Brain Neurips* (Tables S7-S9 and Figure 2A (top)).

With JIND-Multi this reduction in rejection rate is evident even in datasets with minimal batch effect, such as *NSCLC Lung*, where adding batches with JIND already lowers the proportion of rejected cells, but JIND-Multi reduces it even further while achieving similar accuracies (Table S8). In datasets with a considerable batch effect, such as the *Pancreas* dataset, the inclusion of just one additional batch decreases the rejection rate from 9.10% to 4.3% with JIND-Multi, while also improving the accuracy from about 95% to about 97% (Table S7). In contrast, applying JIND under the same conditions exacerbates the rejection rate, increasing it to nearly 20% (Table S7).

In summary, adding more intermediate batches to train JIND-Multi consistently results in a reduction of the rejection rate while maintaining a similar or higher effective accuracy, leading to a higher number of successfully annotated cells (Figure 2 (top)). The decrease in the rejection rate is more pronounced as we add the first additional batches. As we continue to add more intermediate batches this improvement is stabilized. Conversely, in JIND, such enhancement is absent and could potentially elevate the rejection rate, as observed in the *Pancreas* and *Brain Neurips* datasets (Tables S7,S9). In contrast, MARS has not generally shown improved accuracy from the addition of intermediate batches. While in the *Pancreas* dataset the inclusion of two intermediate batches raised mean accuracy from 70% to over 81% (see Table S7), demonstrating this method’s potential in batch effect scenarios, accuracy in the *NSCLC Lung* and *Brain Neurips* datasets declined, with variance even increasing (see Table S8-S9).

Next, we analyzed the prediction accuracy across cell-types. For each dataset, we identify the combination of source-intermediate batches that yielded the best results for MARS on the target data. For these selected experiments, we display the confusion matrix of the best trial from each method (out of 10 runs) and analyze the results. JIND-Multi excels at predicting less-represented cell-types in the source batch by leveraging intermediate batches. In contrast, MARS struggles with imbalances in source cell-types, making it harder to identify less-represented cells. For well-represented cell-types, however, all three methods maintain high accuracy, achieving around 90% (see Figure 3A and Figures S8-S9).

**Figure 3.**
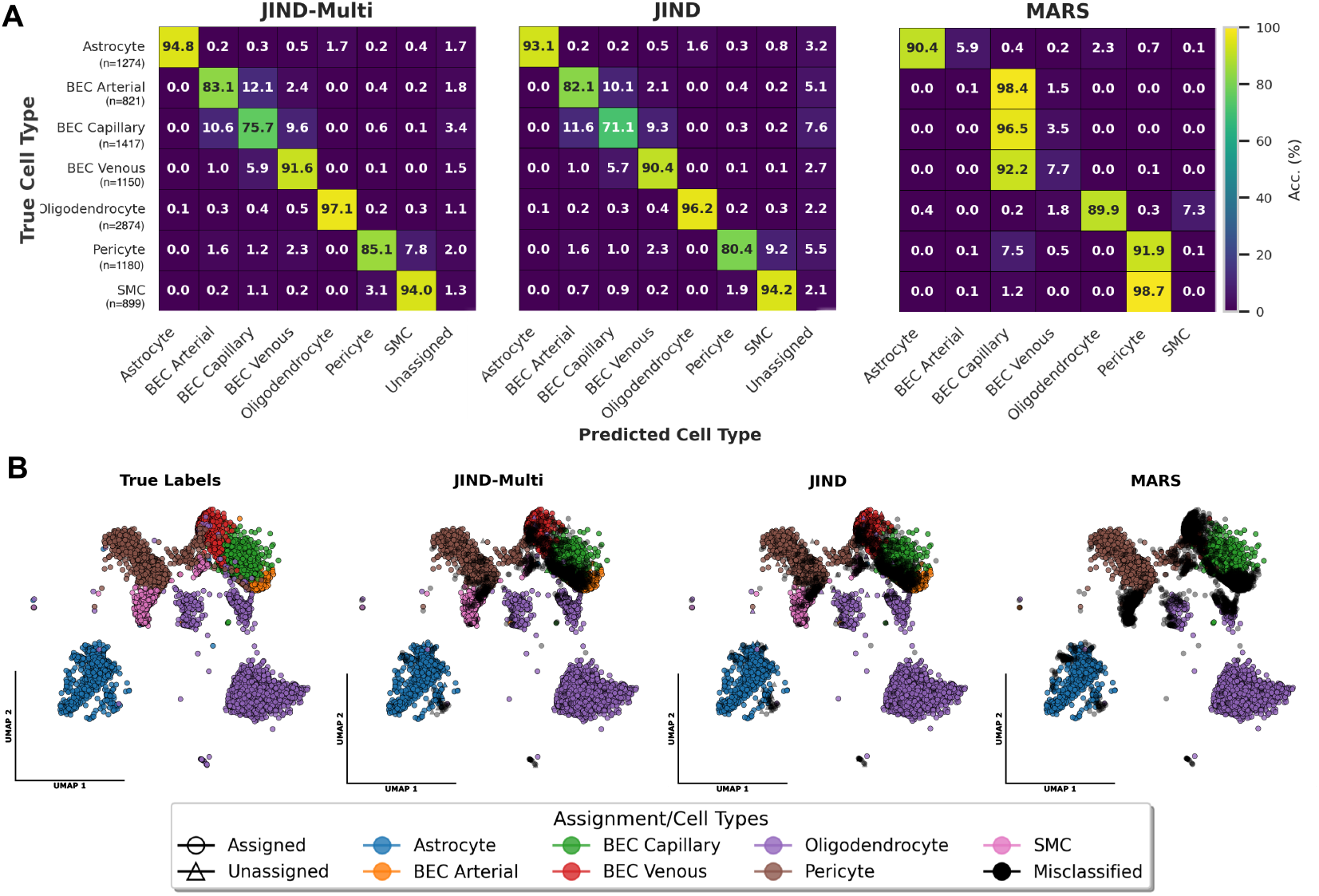
**A**. Confusion matrices showing the prediction accuracies for the different cell-types on the target batch in *Brain Neurips scRNA-Seq* dataset, with results from JIND-Multi trained on batches *C4-AD2* and MARS and JIND trained on batch *C4* with the best trial. JIND-Multi outperforms its predecessor JIND by significantly reducing the number of rejected cells and improving accuracy across all cell-types. However, MARS struggles to differentiate between various types of blood vascular endothelial cells (BEC), as well as between Pericytes and Smooth muscle cells (SMC), which significantly impacts overall accuracy. **B**. UMAP of the cells’ gene expression profiles on the target batch for the *Brain Neurips scRNA-Seq* dataset colored by true labels and predictions got with JIND-Multi, JIND and MARS. For the UMAPs colored by the different method results cells are displayed as circles for assigned predictions in JIND-Multi and JIND, colored by cell-type for correct predictions or in black for misclassified cells. Unassigned cells in JIND-Multi and JIND are denoted by a triangle. The findings underscore MARS’s challenges in discerning between BEC subclasses, its failures in SMC and numerous oligodendrocytes, and its struggles with even more distinct cell-types like astrocytes. Similarly, JIND-Multi encounters complexities in distinguishing between closely related cell-types and subtypes, leading to the classification of many as unassigned.

In the *Pancreas* dataset, the less-represented cell-types, such as *gamma* and *delta* cells, account for just the 3.2% and 7.7% of the source batch, respectively (see Table S1). Here, JIND-Multi demonstrates high precision, achieving better predictions than JIND. For *delta* cells, accuracy increases from 80% in JIND to 88.9% in JIND-Multi, and for *gamma* cells, from 72.2% to 83.3%. MARS, however, completely fails to predict these two cell-types (see Figure S8 and Figure S10), misclassifying them as more abundant celltypes in the source batch, such as *beta* and *alpha* cells. A differential expression (DE) analysis between these cell types revealed at least 1,068 differentially expressed genes between delta and beta cells, and 698 genes between gamma and alpha cells. This reveals a notable misclassification by MARS in this context (see Figure S17-S18).

It should be noted that JIND-Multi and MARS are trained with batches *0, 1, 2*, while JIND is trained solely with batch *0*. For well-represented cell-types (*alpha* and *beta* cells), JIND-Multi improves predictions, reaching accuracies of 94.2% and 96.4%, respectively. Predicting *acinar* cells, representing only 1.4% of the 430 cells in the target batch, remains a challenge for all methods (see Table S1). Only 16.7% of *acinar* cells are correctly predicted in JIND, while JIND-Multi and MARS correctly predict 50%. However, JIND-Multi misclassifies 16.7% of the cells through its filtering scheme, designating the rest as ‘unassigned,’ while MARS misclassifies the remaining 50% entirely.

Globally, among the 27 misclassifications made by JIND-Multi, more than half (51.85%) are labeled as “unassigned.” This filtering mechanism reduces the number of incorrect predictions at the cost of rejecting certain cells, which minimizes its impact on down-stream analyses. By contrast, MARS produces twice as many errors, misclassifying 53 out of the 430 cells in the dataset.

In the *NSCLC Lung* dataset, JIND-Multi and MARS achieve high accuracy for the most represented cell-types, like *B cells* and *monocyte-derived macrophages* using batches *D5-D0-D1-D3-D4-D6* (using 6 batches to train) (see Figure S9 and Figure S11). MARS, however, struggles to distinguish between *CD8* and *CD4 T* cells, with only 88% accuracy.JIND and JIND-Multi surpass 93% in accuracy for these types (see Figure S9). Note however the mean accuracy for this dataset across the 10 runs with MARS was 70.56 *±* 13.12%. see Figure 2A (top) and Table S8.

In the *Brain Neurips* dataset, the least represented cell-types in the source are the Brain Endothelial Arterial Cells (*BEC) arterial* and *Smooth muscle cells* (SMC), at 3.4% and 2.9%, respectively. In the target, they remain the least represented too, at 8.5% and 9.3% (see Table S3). MARS struggles with these cell-types, misclassifying them as the more common *BEC capillary* and *Pericyte* (see Figure 3A-B), which represent the 24.2% and 32.6% of the source. By contrast, JIND-Multi, trained on batches *C4-AD2*, accurately predicts *BEC* subtypes, with accuracies of 83.1% for *BEC arterial*, 75.7% for *BEC capillary*, and 91.6% for *BEC venous*.Differential expression (DE) analysis between the different *BEC* subgroups reveals that misclassifying *BEC arterial* as *BEC capillary* is a much less severe error, with only 237 genes differentially expressed between these two cell-types. In contrast, confusing *BEC venous* with either *BEC arterial* or *BEC capillary*, or misclassifying *Pericyte* as *SMC* cells, represents a more significant error, with 902, 868, and 3210 differentially expressed genes, respectively (see Figure S19-S20).

### Annotation of scATAC-Seq data

To further explore the capabilities of JIND Multi, we explored the ability to reliably annotate on scATAC-Seq data. Analogously to the results obtained in scRNA-Seq data, in scATAC-Seq data JIND-Multi exhibits similar effective accuracies with respect to JIND, while substantially reducing the proportion of rejected cells (Figure 2A (bottom)). For example, the addition of just one extra batch in the *Fetal Kidney* dataset reduces the rejection rate from 5.7% to 1.11%, with higher raw and effective accuracies (Table S12).

A similar trend is observed when more batches are added to JIND, i.e., the rejection rate is reduced and the accuracies maintained. Nevertheless, there is a notable reduction of rejected cells when using JIND-Multi instead of JIND, with values around or below 1% for all cases Figure 2A (bottom). Regarding accuracy values, we note that these are low with respect to scRNA-Seq, with values in the range 70-90% with JIND and JIND-Multi.

With respect to AtacAnnoR, the performance of both methods was found to be similar. JIND achieved a raw accuracy of 89.43% in the evaluated *BMMC* dataset, slightly higher than AtacAnnoR’s accuracy of 89.11% (Table S10).

Focusing on the results by cell-type, we observed that JIND-Multi outperforms JIND by incorporating additional intermediate datasets. Specifically, when training with batches *s4d8-s1d1-s1d2-s1d3-s2d1-s2d4-s2d5-s3d10-s4d1*, important improvements were achieved for various cell-types. *CD8+ T cells*, the most represented cell-type in the source dataset (constituting 53.8% of the 7,601 cells in this batch (Table S4), showed a reduction in rejection rate from 12.4% to 0.5% and an increase in accuracy from 79.4% to 90.2% (see Figure S12). Erythroblasts, which were the least represented cell-type in both the source (3.5% of the batch, see Table S4) and the target dataset (8.8%), also benefited from the addition of cells from batches where they are more prevalent. This enhancement led to an accuracy increase from 85% to 91.9%, a reduction in misclassification as proerythroblasts (a cell-type that has 2,573 DE genes in comparation - see Figure S21), from 11.8% to 6.1%, and a decrease in rejection rate from 2.8% to 0.4%. Additionally, Naive CD20+ cells showed substantial improvements, greatly reducing the rejection rate from 19.3% to 3.8% and the accuracy increasing from 79.1% to 93.6% (see Figure S12-S13).

Compared to AtacAnnoR, the results are slightly superior for some cell-types. It is worth noting that the AtacAnnoR model was trained using scRNA-seq data from the source batch, whereas JIND and JIND-Multi were trained using only the scATAC-seq data. *CD4+ T activated* cells represent a particularly challenging case for JIND, JIND-Multi, and even AtacAnnoR. With the latter, only 74.1% of these cells are correctly classified, while 17.5% are misclassified as *CD8+ T* cells, likely due to the similarity in their expression profiles with 519 DE genes (see Figure S12,S21). In this scenario, the reduction in the rejection rate achieved by JIND-Multi has unfortunately increased the misclassification error, with the proportion of CD4+ T activated cells incorrectly predicted as CD8+ T increasing from 46.5% to 60.5% (Figure S12).

### Model robustness

To evaluate the robustness of JIND-Multi across different datasets, we analyzed its performance and variability in predictions over 10 trials, comparing it to other methods such as MARS (for scRNA-Seq) and AtacAnnoR (for scATAC-Seq). Specifically, we assessed how many cells were predicted incorrectly across different ranges of trials using the same combination of source-intermediate batches that yielded the best results for MARS on the target data.

For scRNA-Seq data, JIND-Multi demonstrated greater robustness overall, with most of its misclassified cells also being misclassified by MARS (see Figure 2B, Figure 3, Figure S10-S11,S13). Although both JIND-Multi and MARS exhibit some persistent errors across all the trials, MARS shows more unstable and variable behavior, with higher frequencies of misclassification at all trial ranges. For instance, in *NSCLC Lung* dataset, JIND-Multi misclassifies 15 cells (1.76% of the whole target batch) in 7-9 trials, 7 cells (0.8%) in 4-6 trials, and 4 cells (0.5%) in 1-3 trials. By contrast, MARS misclassified 30 cells (3.52%) in 7-9 trials, 319 cells (37.4%) in 4-6 trials, and 309 cells (36.2%) in 1-3 trials (see Figure 2B). A similar trend is observed in the *Brain Neurips* dataset.

For scATAC-Seq data, robustness was analyzed in *BMMCscATAC-Seq* across batches *s4d8-s1d1-s1d2-s1d3-s2d1-s2d4-s2d5-s3d10-s4d1* for JIND-Multi and *s4d8* for AtacAnnoR. JIND-Multi consistently failed to predict 218 cells across all 10 trials, compared to 112 for AtacAnnoR. However, for other ranges of trial counts, AtacAnnoR showed higher numbers of misclassified cells. This suggests that even though JIND-Multi is unable to predict a smaller sample of cells reliably, it retains higher consistency in its predictions overall (see Figure 2B).

In summary, JIND-Multi demonstrated greater robustness on both scRNA-Seq and scATAC-Seq datasets, with fewer inconsistencies in predictions compared to MARS and AtacAnnoR, making it a more reliable choice for accurate cell-type annotations.

### Running time

JIND-Multi exhibits longer execution times, as expected, as it handles more batches than its predecessor (Tables S13-S18). However, this time depends on the amount of samples and batches available. For example, in the *NSCLC Lung* scRNA-Seq dataset, JIND took 50 seconds to execute regardless of whether it used one batch or multiple batches, while JIND-Multi took approximately 750 seconds when processing the 6 batches.

In scATAC-Seq, AtacAnnoR is more time-efficient, requiring approximately half the execution time of JIND when processing a single batch (84 seconds for JIND vs. 44 seconds for AtacAnnoR) in *BMMC scATAC-Seq* dataset (Table S16). With JIND-Multi, processing additional batches increases the execution time, reaching 169 seconds for two batches and further increasing with more intermediate batches.

## Conclusions

In this work we presented JIND-Multi, a tool for the automatic annotation of single-cell RNA and ATAC data. JIND-Multi extends upon JIND, supporting the use of several annotated datasets for label transfer. Our experimental findings demonstrate that JIND-Multi effectively integrates diverse annotated batches, yielding high-confidence predictions and mitigating rejection rates for both scRNA-Seq and scATAC-Seq data. An additional advantage of JIND-Multi is that it does not require paired scRNA-Seq data for the annotation of scATAC-Seq. These attributes position JIND-Multi as a potentially valuable asset for single-cell atlas annotation and atlas generation or evaluation.

## Methods

As mentioned above, several established methods were also utilized in this work. The corresponding code and data for these methods can be accessed at the following links:

- **AtacAnnoR**: https://github.com/TianLab-Bioinfo/AtacAnnoR
- **MARS**: http://snap.stanford.edu/mars/
- **JIND**: https://github.com/mohit1997/JIND

## Availability of source code and requirements

Lists the following:

- Project name: JIND-Multi
- Project home page: https://github.com/ML4BM-Lab/JIND-Multi/releases/tag/v1.0.0
- Operating system(s): e.g. Platform independent
- Programming language: Python
- Other requirements: Python 3.6.8 or higher, miniconda.
- License: MIT License

## Data availability

The datasets supporting the results of this article are available from the Zenodo repository [23].

## Abbreviations

scRNA-Seq: Single-Cell RNA Sequencing; scATAC-Seq: Single-Cell ATAC Sequencing; NN: Neural Networks; GAN: Generative Adversarial Network; NSCLC: Non-Small Cell Lung Cancer; BMMC: Human Bone Marrow Mononuclear Cells.

## Author’s Contributions

J.S., and A.K.wrote the JIND-Multi software with O.G. supporting; M.H., G.S., and I.O. provided supervision. J.S. developed the GitHub repository for the project with X.G. providing additional support.

J.S. wrote the original draft of the manuscript with all authors reviewing.

