## Supplementary material for "JIND-Multi: Leveraging Multiple Labeled Datasets for Automated Annotation of Single-Cell RNA and ATAC Data": JIND_Multi_supplementary.pdf

### Supplementary data for manuscript "Jind-Multi: An extended framework to leverage multiple labeled datasets for the automatic annotation of single-cell RNA and ATAC data"

Joseba Sancho[1], Akash Kanhirodan[2], Xabier Garrote[1], Olivier Gevaert[3], Mikel Hernaez[4,5], Guillermo Serrano[1,6], and Idoia Ochoa[1,5]

[1] *Tecnun School of Engineering, Universidad de Navarra, Donostia, Spain.*
[2] *National Institute of Technology Calicut, India.*
[3] *Stanford Center for Biomedical Informatics Research, Stanford University, California.*
[4] *Centro de Invesstigacion Medica Aplicada (CIMA), Universidad de Navarra, Spain.*
[5] *Instituto de Ciencia de los Datos e Inteligencia Artificial (DATAI), Universidad de Navarra, Spain.*
[6] *Biological and Environmental Science and Engineering Division, King Abdullah University of Science and Technology (KAUST), Thuwal, Saudi Arabia.*

### 1 Findings

JIND-Multi consists of three steps (see Figure S1). In brief, the first step trains the encoder-classifier with the source batch. The second step aligns the additional labeled datasets to the source batch, and finally the third step performs the cell-type inference on the unlabeled dataset. The overall architecture is equivalent to that of JIND [3] (see Figure S1, Step 1), with the novelty coming on the second step, where more annotated datasets are incorporated into the model for improved inference (see Figure S1, Step 2). Next, we describe these steps in more detail.

#### 1.1 Prediction model (Step 1)

The prediction model in the first step is composed of two subnetworks: an encoder and a classifier. Let $x, y$ be the expression data of a cell containing $M$ genes (the $5,000$ genes with the highest variance by default) and the corresponding cell-type, respectively. The encoder takes $x$ as input and generates a latent space $h$ (colored green in Figure S1, Step 1) of reduced dimension that serves as input to the classifier. The latter computes the predictions, a K-dimensional vector $\hat{y}$ that represents the probability of the cell belonging to each of the $K$ classes. The architecture applied in this step for both encoder and classifier is the same as in JIND's [3], using ReLU (Rectified Linear Unit) non-linearity as the activation function and a latent space of dimension 256.

**Training:** The prediction model is trained using one of the labeled datasets, denoted as the source batch $(X^s, Y^s)$ (see Figure S1, Step 1). The source dataset is split into training (80%) and validation sets, performing a stratified partitioning of the samples based on cell-types. The parameters of the model are trained by minimizing the weighted categorical cross-entropy loss $\mathcal{L}_C(y, \hat{y})$. Weights are added to the loss to account for possible class imbalance in the data. See [3] for more details. The training is carried out using a batch size of 128 and 15 epochs. We use the Adam optimizer with a learning rate of $10^{-3}$ that decreases as training progresses.

**Source batch selection:** The user can specify the annotated batch to be used as source as an input parameter to JIND-Multi. We recommend using the one with the highest number of cells for each cell-type. Alternatively, if no batch is specified, JIND-Multi will select as source the batch that produces the least amount of rejected cells on the target batch when used as source in JIND (i.e., without additional intermediate batches).

**Filtering:** JIND-Multi employs $K$ confidence thresholds to filter cell predictions based on cell-type, ensuring robustness against misclassifications during inference, as detailed in the previous method [3]. This filtering mechanism is integrated into the training of the prediction model utilizing for calculation samples from the validation set. The thresholds calculated in this step remain fixed and are not updated in subsequent steps of the pipeline. When predicting on an unlabeled (target) batch, samples for which the highest probability in the prediction do not exceed the threshold for the corresponding cell-type are marked as "Unassigned".

#### 1.2 Adapt model (Step 2)

Once the encoder and classifier are trained, the remaining available annotated batches are processed by JIND-Multi one by one (see Figure S1, Step 2). Let $x \in \mathcal{R}^M$ be a sample from an intermediate batch $j$ $(X^j)$. Passing the data through the encoder trained in the previous step will generate a latent code $h$ (represented in red in Figure S1) not aligned with the latent space of the source batch. This misalignment is due to the potential presence of batch effects between the source and intermediate batches.

To align the distribution of the intermediate batch with that of the source, we train a generator $G^j$ that shifts and scales $h$ to obtain a modified latent space $\hat{h}$ that can be used by the already trained classifier. This generator consists of two Neural Network (NN) blocks $S^j$ and $B^j$ with the same architecture, consisting as in JIND [3] of two fully connected layers with 512 neurons each (followed by a ReLU activation function), and a fully connected layer that generates the 256-dimensional $\hat{h}$ output. Both $S^j$ and $B^j$ take the cell's gene expression vector $x$ as input and generate a corrected latent code $\hat{h}$ using a mapping based on residual connections, determining the necessary scale and bias adjustments to alter the latent code [4]. This can be formulated as:

$$\hat{h} = G^j(x, h) = S^j(x) \circ h + B^j(x), \tag{1}$$

where $\circ$ indicates element-wise multiplication.

**Training:** Given that we already have the cell-types $Y^j$, the parameters of $S^j$ and $B^j$ are optimized by again minimizing the weighted cross-entropy loss $\mathcal{L}_C(y, \hat{y})$, while keeping the parameters of the encoder and classifier fixed. The generator's parameters, as in JIND [3], are set up so that initially $G^j$ acts as an identity mapping, resulting in $\hat{h} = h$. The adapt model is trained with the following configurations: 15 epochs, a learning rate of $10^{-4}$ and a batch size of 128. Additionally, the performance of this integration is validated using 10% of the intermediate set. Note that the employed learning rate scheduler is the same as that used in the classifier training.

##### 1.2.1 Fine-tuning the encoder and classifier

After adapting an intermediate set $i$ to the source's latent space, a parallel fine-tuning of the encoder and the classifier is conducted using the already trained sets, i.e., the source batch and intermediate batches $[1 : i]$. The parameters are adjusted by sequentially passing each dataset through its respective model, while keeping the parameters of the adapt models $S^j$ and $B^j$ fixed, for $j \in [1 : i]$. Note therefore that this fine-tuning is performed $n$ times, with $n$ being the number of intermediate batches.

#### 1.3 Final Model (Step 3)

##### 1.3.1 Inference Stage (Step 3.A)

Once the training is finalized (Steps 1 and 2), JIND-Multi can be employed to annotate a new batch, referred to as the target batch $X^t$, for which cell annotations are not available. To align the latent space $h$ generated by passing the input data $x \in X^t$ through the trained encoder, a Generative Adversarial Network (GAN) is trained, similarly as implemented in JIND [3] (see Figure S1, Step 3.A). The GAN consists of a discriminator $D$ and a generator $G^t$. $D$ is a NN classifier whose goal is to distinguish between input data originating from the latent code of the sources (source and intermediate batches) and input data originating from the target. On the other hand, the task of the generator is to deceive the discriminator by creating a latent code $\hat{h}$ that is indistinguishable from the latent code generated from the source batches. $G^t$ uses the same architecture as the generators used for the intermediate batches, and $D$ is based on fully connected layers with a single output neuron with sigmoid activation function (refer to [3] for additional details).

**Training:** $D$ and $G^t$ are trained competitively, jointly minimizing the losses $\mathcal{L}_D$ and $\mathcal{L}_G$ of both networks as described in detail in [3]. The parameters of $D$ are optimized through minimizing the loss $\mathcal{L}_D$, while the parameters $S^t$ and $B^t$ of $G^t$ are optimized through minimizing $\mathcal{L}_G$. To prevent overfitting of $G^t$, the objective function is regularized by heavily penalizing the L2-norm of its parameters, as performed in JIND [3].

All annotated samples are used for training the GAN. Specifically, for each epoch, samples from the target batch's latent code (referred to as 'negative' examples) are sequentially passed through $D$ along with samples from any of the annotated batch's latent code ('positive' examples) through their corresponding model (refer to Figure S1, Step 3.A). Note that during the GAN training, the parameters of the encoder, $S^i_j$ and $B^i_j$, for $j \in [1 : n]$, remain fixed.

Both the $G^t$ and $D$ are trained using the RMSprop optimizer with a learning rate of $10^{-4}$. The $G^t$ employs a weight decay of $10^{-2}$, while the $D$ uses $10^{-3}$. Gaussian noise, with a default sigma value of zero, is introduced during training. Training is conducted with a batch size of 128 over 15 epochs. Early stopping is implemented, halting training if there's no improvement in loss for seven consecutive epochs.

With JIND-Multi, $D$ benefits from access to a larger pool of samples, thereby increasing the variability it captures. This enhanced variability improves $D$'s ability to provide more informative feedback to $G^t$. As a result, $G^t$ can generate a latent space that produces better predictions when inputted into the classifier.

##### 1.3.2 Final prediction model (Step 3.B)

Once the generator $G^t$ for the target batch has been trained through adversarial training, the final prediction model is ready to make cell-type predictions on the target batch (Figure S1, Step 3.B). A cell's gene expression vector $x$ passes through the trained encoder and generator, creating the corrected latent code, which undergoes ReLU activation before being input into the classifier to calculate the inferred K-probabilities of the cell belonging to each of the considered cell-types.

An additional fine-tuning step is performed, as in JIND+ [3], on the encoder's and the classifier's parameters using the cells (samples) from the target batch that have obtained high confidence in the prediction. The resulting model is then used to compute the final predictions. Specifically, for each cell, JIND-Multi outputs the class with the highest probability, as long as this probability surpasses the threshold determined for that cell type during training. Otherwise, the 'Unassigned' label is set.

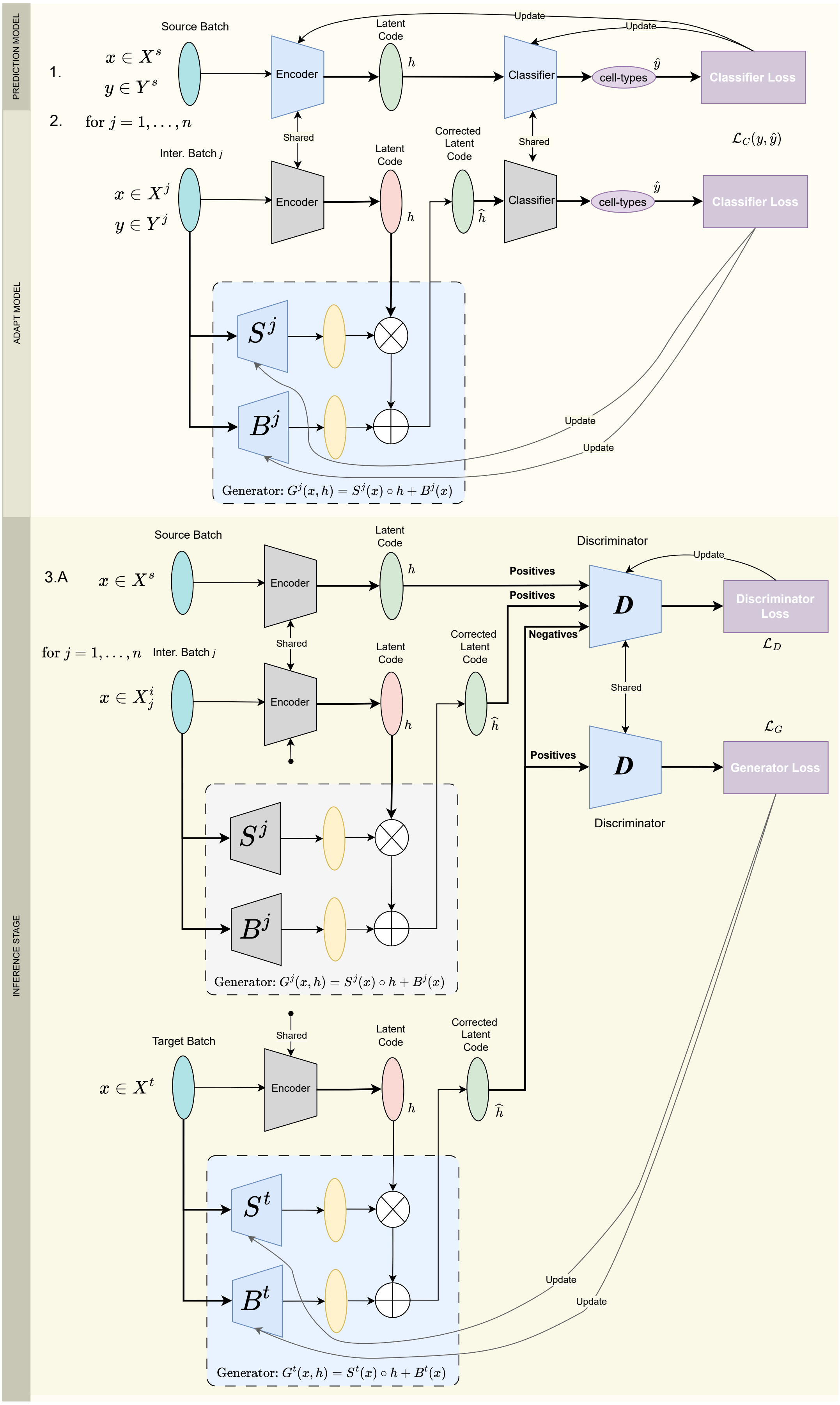
PREDICTION MODEL
ADAPT MODEL
INFERENCE STAGE
Source Batch
1.
$x \in X^s$
$y \in Y^s$
Encoder
Latent Code
$h$
Classifier
cell-types
$\hat{y}$
Classifier Loss
Update
Update
2.
for $j = 1, \ldots, n$
Shared
Shared
Inter. Batch $j$
$x \in X^j$
$y \in Y^j$
Encoder
Latent Code
Corrected Latent Code
$\hat{h}$
Classifier
cell-types
Classifier Loss
$\mathcal{L}_C(y, \hat{y})$
$S^j$
$B^j$
Update
Update
Generator: $G^j(x,h) = S^j(x) \circ h + B^j(x)$
Source Batch
3.A
$x \in X^s$
Encoder
Latent Code
Discriminator
Positives
Positives
Negatives
$D$
Discriminator Loss
$\mathcal{L}_D$
for $j = 1, \ldots, n$
Inter. Batch $j$
$x \in X^i_j$
Shared
Corrected Latent Code
$\mathcal{L}_G$
Positives
$D$
Generator Loss
Discriminator
$S^j$
$B^j$
Generator: $G^j(x,h) = S^j(x) \circ h + B^j(x)$
Target Batch
$x \in X^t$
Encoder
Latent Code
Corrected Latent Code
$S^t$
$B^t$
Update
Update
Generator: $G^t(x,h) = S^t(x) \circ h + B^t(x)$

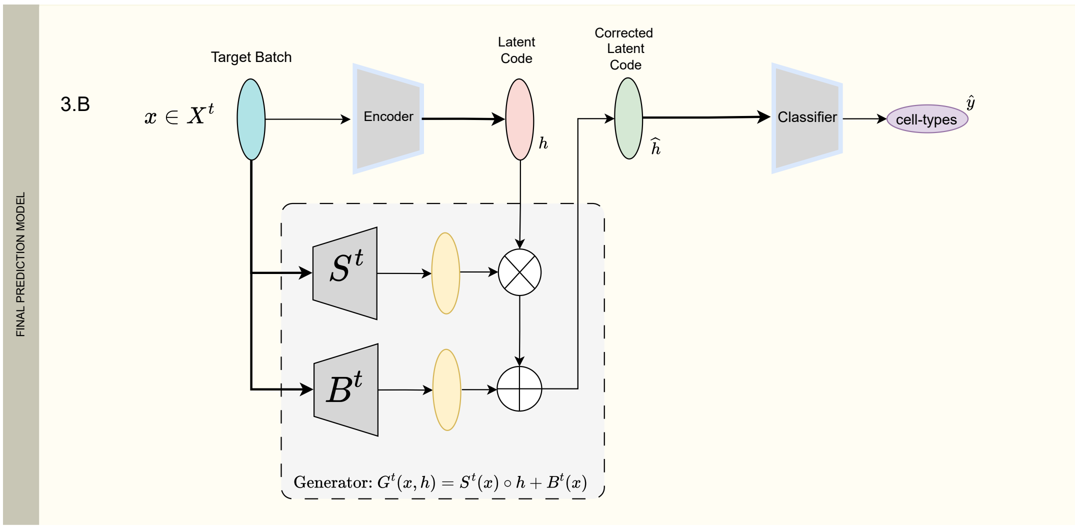

Figure 1: Schematic of JIND-Multi. *Prediction Model* (**Step 1**): The encoder and classifier are trained using the source batch. A latent code (depicted in green) is generated from the source batch, enabling cell annotation when applied to the classifier. *Adapt Model* (**Step 2**): A generator is trained to mitigate batch effects from each intermediate set and align their latent code (depicted in red) with that of the source (depicted in green after going through the trained generator). Parameters optimized during training are shown in light blue, while fixed parameters are depicted in gray. *Inference Stage on an unlabeled batch* (**Step 3.A**): A Generative Adversarial Network (GAN) is employed to align the target's latent code with that of the sources. *Final Prediction Model* (**Step 3.B**): The target batch is processed through the encoder and the generator trained in Step 3.A to produce a latent code capable of predicting cell types when input to the classifier. The encoder and classifier undergo a final fine-tuning (indicated by light blue underline) using the highest-confidence predictions on the target batch.

### 2 Data

#### 2.1 Data Preprocessing

**scRNA-Seq:** Data was processed using Scanpy (v1.9.5). The preprocessing consisted of removing the cells with less than 2500 features with at least one count per cell and a mitochondrial content higher than 5%. Data was normalized and scaled regressing out the data using as covariates the total number of counts to mitigate the technical sequencing depth from batch effect.

**scATAC-Seq:** Data was preprocessed using EpiScanpy (v0.4.0). The identity of the peaks was assigned by proximity to the transcription start site for each feature. Similarly to the scRNA-Seq counterpart, we kept the cells expressing more than 100000 features and removed the cells with more than 5% of the total of fragments assigned to the mitochondrial genes. Finally, data was normalized regressing as covariate the total number of counts per cell and scaled.

For both scRNA-Seq and scATAC-Seq data, we select the top $5,000$ genes with the highest variance across batches from the batched data and filter the cells based on a minimum cell count per batch. The set number depends on the experiment under consideration.

#### 2.2 Datasets

In this study, we consider three scRNA-Seq datasets denoted as *Pancreas*, *NSCLC Lung*, and *Brain Neurips*, and three scATAC-Seq datasets denoted as *BMMC*, *Fetal Heart*, and *Fetal Kidney*.

All datasets can be downloaded from the following link: `https://doi.org/10.5281/zenodo.14000644`.

***Pancreas* scRNA-Seq dataset:** It comprises 4 batches, 6 cell-types (acinar, alpha, beta, delta, ductal, and gamma cells), 2,448 genes, and approximately 12 thousand cells. This dataset was obtained after applying a filtering criterion to retain only those cell-types with at least 5 cells per batch. See Table S1 for the resulting cell-type proportions and number of cells per batch.

| Batch Name | acinar | alpha | beta | delta | ductal | gamma | Total cells |
|---|---|---|---|---|---|---|---|
| 0 (source) | 12.3% | 30.0% | 32.6% | 7.7% | 13.9% | 3.2% | 7742 |
| 1 | 10.8% | 40.2% | 22.2% | 9.5% | 12.1% | 5.0% | 2018 |
| 2 | 9.0% | 43.4% | 13.2% | 5.5% | 18.9% | 9.6% | 2038 |
| 3 (target) | 1.4% | 44.1% | 25.8% | 2.0% | 22.3% | 4.1% | 430 |
| Total | | | | | | | 12228 |

Table 1: Pancreas scRNA-Seq dataset description after cell filtering. Source and target batches used for the experiments are specified.

***NSCLC Lung* scRNA-Seq dataset:** This dataset comprises more than 6 thousand cells distributed in 7 batches, 3,448 genes, and 4 cell-types (B-Cells, CD4 T-Cells, CD8 T-Cells, and monocyte-derived macrophages (MDMs)) (Table S2). The cells were filtered with a minimum of 20 cells per cell-type for every batch.

| Batch Name | B cells | CD4 T cells | CD8 T cells | Monocyte-derived Mph | Total cells |
|---|---|---|---|---|---|
| Donor 0 | 2.4% | 35.4% | 55.2% | 6.8% | 806 |
| Donor 1 | 11.4% | 53.1% | 25.3% | 10.1% | 822 |
| Donor 2 (target) | 9.2% | 45.4% | 35.0% | 10.2% | 853 |
| Donor 3 | 50.6% | 25.5% | 19.0% | 4.7% | 715 |
| Donor 4 | 3.9% | 53.6% | 19.4% | 22.9% | 1080 |
| Donor 5 (source) | 13.1% | 23.7% | 26.0% | 37.0% | 1702 |
| Donor 6 | 20.6% | 39.2% | 13.0% | 27.0% | 499 |
| Total | | | | | 6477 |

Table 2: NSCLC Lung scRNA-Seq dataset description after cell filtering. Source and target batches used for the experiments are specified.

***Brain Neurips* scRNA-Seq dataset:** This dataset has an immense collection of cells and batches, and we selected for the analysis the 6 batches with the highest number of cells, compromising in total 95 thousand cells distributed among 7 cell-types (astrocyte, BEC arterial, BEC capillary, BEC venous, oligodendrocyte, pericyte and vascular smooth muscle cells (SMC)) (Table S3). Batches were filtered to retain only those cell-types with at least 100 cells per cell-type for each batch.

| Batch Name | astrocyte | BEC arterial | BEC capillary | BEC Venous | oligodendrocyte | pericyte | SMC | Total cells |
|---|---|---|---|---|---|---|---|---|
| C4 (source) | 13.5% | 3.4% | 24.2% | 10.6% | 12.4% | 32.6% | 2.9% | 20566 |
| ADx2 | 17.2% | 13.3% | 3.7% | 3.2% | 42.7% | 18.7% | 0.9% | 15130 |
| C7 (target) | 13.2% | 8.5% | 14.7% | 11.9% | 29.8% | 12.2% | 9.3% | 9615 |
| ADx1 | 17.2% | 13.2% | 3.8% | 3.2% | 42.7% | 18.7% | 0.9% | 15131 |
| ADx4 | 19.2% | 12.9% | 3.4% | 1.5% | 45.8% | 15.7% | 1.4% | 18388 |
| AD2 | 10.9% | 8.3% | 22.2% | 7.4% | 14.0% | 30.6% | 6.2% | 17009 |
| Total | | | | | | | | 95839 |

Table 3: Brain Neurips scRNA-Seq dataset description after cell filtering. Source and target batches used for the experiments are specified.

***BMMC* scATAC-Seq dataset:** We selected 10 batches from the *BMMC* dataset, comprising a total of 41,953 cells across 7 cell-types (CD14+ Mono, CD4+ T activated, CD8+ T, erythroblast, NK, naive CD20+ B, and proerythroblast) (Table S4). The minimum number of cells per cell-type was set to 18.

| Batch Name | CD14+ Mono | CD4+T activated | CD8+T | erythroblast | NK | Naive CD20+ B | proerythroblast | Total cells |
|---|---|---|---|---|---|---|---|---|
| s2d4 | 26.6% | 8.9% | 5.3% | 34.6% | 3.1% | 13.7% | 7.3% | 4282 |
| s3d3 (target) | 30.8% | 8.2% | 13.6% | 8.8% | 17.2% | 14.1% | 7.1% | 2774 |
| s4d8 (source) | 6.0% | 9.0% | 53.8% | 3.5% | 5.5% | 11.6% | 10.5% | 7601 |
| s1d3 | 8.1% | 13.1% | 34.3% | 12.5% | 23.4% | 4.4% | 3.9% | 2973 |
| s4d1 | 20.5% | 12.5% | 15.3% | 9.6% | 22.7% | 12.6% | 6.5% | 4938 |
| s2d1 | 42.9% | 14.6% | 5.5% | 15.7% | 8.8% | 9.4% | 2.7% | 3044 |
| s1d2 | 16.3% | 16.4% | 37.8% | 3.1% | 22.7% | 2.8% | 0.7% | 4314 |
| s3d10 | 43.2% | 5.5% | 11.6% | 10.5% | 14.6% | 11.3% | 2.9% | 3710 |
| s1d1 | 21.3% | 19.3% | 16.7% | 18.4% | 13.6% | 8.9% | 1.5% | 4321 |
| s2d5 | 42.9% | 12.3% | 18.8% | 1.6% | 21.2% | 2.6% | 0.4% | 3996 |
| Total | | | | | | | | 41953 |

Table 4: BMMC scATAC-Seq dataset description after cell filtering. Source and target batches used for the experiments are specified.

***Fetal Heart* scATAC-Seq dataset:** This dataset comprises 3 batches with more than 30,000 cells across 6 cell-types (fetal atrial cardiomyocyte, fetal cardiac fibroblast, fetal endocardial cells, fetal endothelial (general)1, fetal fibroblast (general)3, and fetal ventricular cardiocyte) and 12,436 genes (Table S5). The minimum number of cells per cell-type was set to 100.

| Batch Name | Atrial Cardiomyocyte | Cardiac Fibroblast | Endocardial Cell | Endothelial (General)1 | Fibroblast (General)3 | Ventricular Cardiocyte | T.cells |
|---|---|---|---|---|---|---|---|
| heart sample 39 (source) | 7.4% | 15.5% | 5.8% | 5.0% | 1.8% | 64.2% | 16200 |
| heart sample 32 | 9.5% | 13.9% | 1.6% | 10.6% | 4.1% | 60.03% | 7733 |
| heart sample 14 (target) | 9.8% | 17.8% | 5.7% | 6.6% | 1.9% | 57.9% | 6457 |
| Total | | | | | | | 30390 |

Table 5: Fetal Heart scATAC-Seq dataset description after cell filtering. Source and target batches used for the experiments are specified.

***Fetal Kidney* scATAC-Seq dataset:** This dataset is composed of 4 batches containing 19,107 cells, and includes 4 cell-types: mesangial cell 1, mesangial cell 2, metanephric cell, and ureteric bud cell (Table S6). We used a minimum of 100 cells for each cell-type.

| Batch Name | Mesangial Cell 1 | Mesangial Cell 2 | Metanephric Cell | Ureteric Bud Cell | Total cells |
|---|---|---|---|---|---|
| kidney sample 34 | 15.9% | 7.0% | 63.0% | 14.1% | 5215 |
| kidney sample 65 | 18.9% | 7.1% | 58.6% | 15.4% | 2331 |
| kidney sample 67 (target) | 16.5% | 7.7% | 64.1% | 11.8% | 1572 |
| kidney sample 3 (source) | 24.1% | 6.2% | 62.9% | 6.8% | 9989 |
| Total | | | | | 19107 |

Table 6: Fetal Kidney scATAC-Seq dataset description after cell filtering. Source and target batches used for the experiments are specified.

#### 2.3 Analysing the presence of batch effect

To analyze potential batch effects within batches of a given dataset, we visualize the gene expression profiles of each cell in a dataset with UMAP [5]. Figures S2-S7 show the results of the UMAP analysis for each considered dataset, with samples from each dataset colored by: i) their cell-type (left); ii) or the batch they belong to (right). This allows us to differentiate inherent biological variations due to the cell-type from the presence of artifacts due to the batch effect. In the absence of batch effect, samples of the same cell-type are expected to occupy the same space in the UMAP for the different batches.

Visualizing the UMAP representation of the datasets reveals a significant batch effect in the *Pancreas* scRNA-Seq dataset (Figure S2). This may be explained by the nature of this dataset, as it comes from three different experiments [1], [6] and [7]. The other datasets show a batch effect to a lesser extent, with the *NSCLC Lung* scRNA-Seq dataset exhibiting the least batch effect.

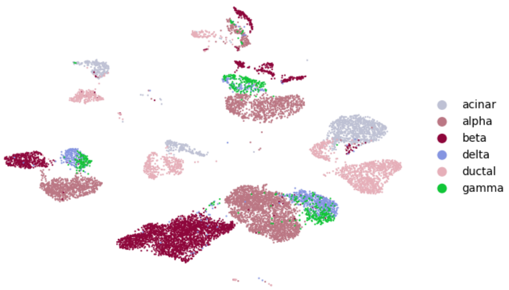

(a) Cell Type

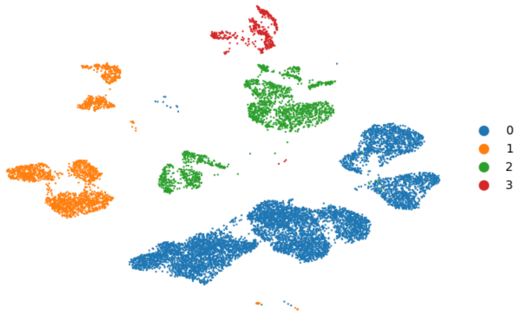

(b) Batch

Figure 2: UMAP of the cells' gene expression profiles of the *Pancreas* scRNA-Seq dataset colored by a) cell type and b) batch.

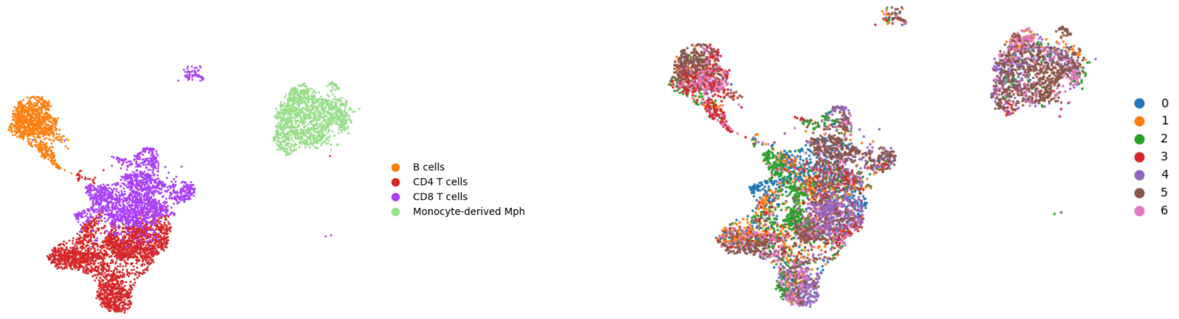

(a) Cell Type

(b) Batch

Figure 3: UMAP of the cells' gene expression profiles of the *NSCLC Lung* scRNA-Seq dataset colored by a) cell type and b) batch.

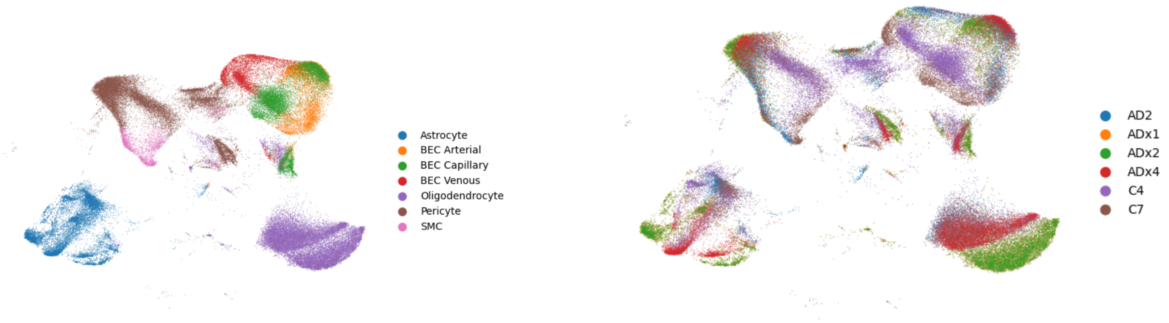

(a) Cell Type

(b) Batch

Figure 4: UMAP of the cells' gene expression profiles of the *Brain Neurips* scRNA-Seq dataset colored by a) cell type and b) batch.

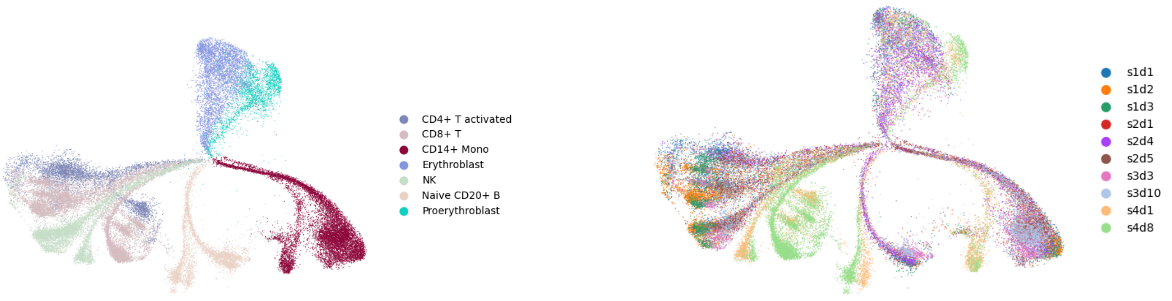

(a) Cell Type

(b) Batch

Figure 5: UMAP of the cells' gene expression profiles of the *BMMC* scATAC-Seq dataset colored by a) cell type and b) batch.

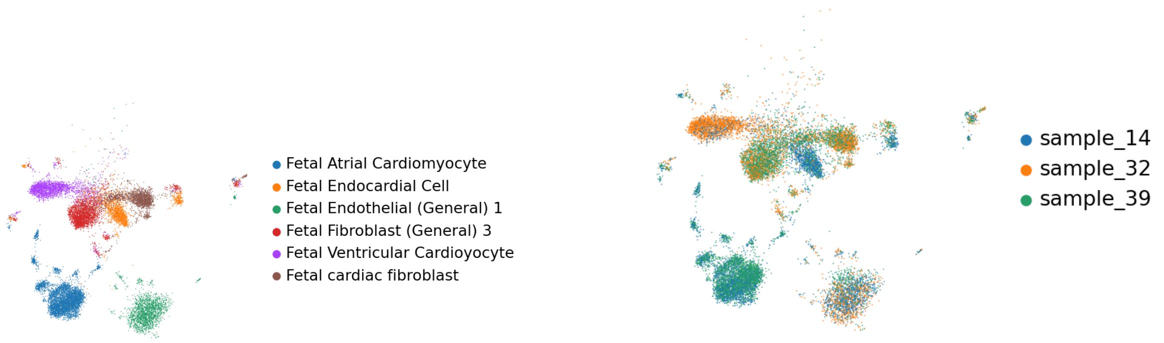

(a) Cell Type (b) Batch

Figure 6: UMAP of the cells' gene expression profiles of the *Fetal Heart* scATAC-Seq dataset colored by a) cell type and b) batch.

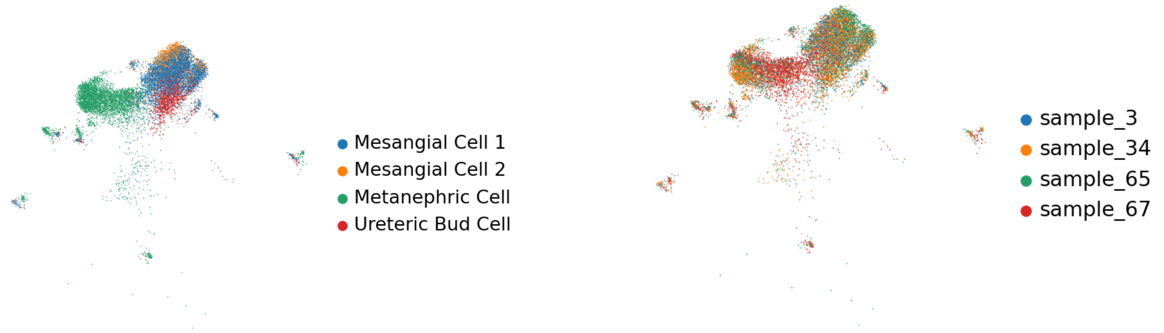

(a) Cell Type (b) Batch

Figure 7: UMAP of the cells' gene expression profiles of the *Fetal kidney* scATAC-Seq dataset colored by a) cell type and b) batch.

### 3 Extended results

#### 3.1 scRNA-Seq experiments

For the annotation of scRNA-Seq data, we compare JIND-Multi with JIND and MARS [2], an existing method that can also use multiple labeled batches for training. The results for the three considered scRNA-Seq datasets, across 10 runs, are shown below. Since MARS does not support filtering cells based on the prediction confidence, we compare the raw accuracy between the methods. We observe that in all cases JIND-Multi exhibits a superior performance.

| Batches | JIND-Multi | | | JIND | | | MARS |
|---|---|---|---|---|---|---|---|
| | raw. acc | eff.acc | rej | raw.acc | eff.acc | rej | raw.acc |
| *0 → 3* | | | | 91.62 ± 0.00 | 94.88 ± 0.00 | 9.10 ± 0.00 | 70.06 ± 3.92 |
| *0-1 → 3* | 95.55 ± 0.26 | 96.96 ± 0.24 | 4.32 ± 0.30 | 96.51 ± 0.00 | 99.42 ± 0.00 | 19.10 ± 0.00 | 79.93 ± 5.54 |
| *0-1-2 → 3* | 96.02 ± 0.51 | 97.71 ± 0.78 | 6.08 ± 2.83 | 96.32 ± 0.42 | 98.68 ± 0.16 | 16.45 ± 2.97 | 81.88 ± 4.77 |

Table 7: Performance of JIND-Multi versus JIND and MARS for the *Pancreas* scRNA-Seq dataset. Batches column indicate the source batches (left of arrow) and target batch (right of arrow). Note that JIND-Multi with one labeled batch is equivalent to JIND.

| Batches | JIND-Multi | | | JIND | | | MARS |
|---|---|---|---|---|---|---|---|
| | raw. acc | eff.acc | rej | raw.acc | eff.acc | rej | raw.acc |
| *D5 → D2* | | | | 94.49 ± 0.00 | 96.70 ± 0.00 | 7.50 ± 0.00 | 80.25 ± 11.33 |
| *D5-D0 → D2* | 91.77 ± 0.51 | 92.67 ± 0.41 | 2.70 ± 0.20 | 94.02 ± 0.00 | 94.84 ± 0.00 | 2.20 ± 0.00 | 72.29 ± 10.35 |
| *D5-D0-D1 → D2* | 95.33 ± 0.23 | 96.03 ± 0.27 | 1.72 ± 0.27 | 94.13 ± 0.28 | 95.25 ± 0.13 | 3.16 ± 0.29 | 69.16 ± 10.48 |
| *D5-D0-D1-D3 → D2* | 95.36 ± 0.19 | 96.30 ± 0.11 | 1.73 ± 0.23 | 95.31 ± 0.12 | 96.22 ± 0.15 | 2.17 ± 0.22 | 66.33 ± 9.41 |
| *D5-D0-D1-D3-D4 → D2* | 95.22 ± 0.17 | 95.93 ± 0.08 | 1.13 ± 0.17 | 94.56 ± 0.58 | 95.70 ± 0.38 | 2.59 ± 0.76 | 64.00 ± 8.96 |
| *D5-D0-D1-D3-D4-D6 → D2* | 95.07 ± 0.12 | 95.69 ± 0.16 | 1.17 ± 0.32 | 94.91 ± 0.24 | 95.80 ± 0.28 | 2.43 ± 0.30 | 70.56 ± 13.12 |

Table 8: Performance of JIND-Multi versus JIND and MARS for the *NSCLC Lung* scRNA-Seq dataset. Batches column indicate the source batches (left of arrow) and target batch (right of arrow). For JIND, when more than one labeled batch is used, the batches are merged into one.

| Batches | JIND-Multi | | | JIND | | | MARS |
|---|---|---|---|---|---|---|---|
| | raw. acc | eff.acc | rej | raw.acc | eff.acc | rej | raw.acc |
| *C4 → C7* | | | | 90.35 ± 0.00 | 91.58 ± 0.00 | 3.80 ± 0.00 | 64.17 ± 0.65 |
| *C4-AD2 → C7* | 90.81 ± 0.19 | 91.41 ± 0.27 | 1.68 ± 0.14 | 89.39 ± 0.28 | 90.44 ± 0.26 | 2.80 ± 0.20 | 63.45 ± 0.55 |
| *C4-AD2-ADx1 → C7* | 90.52 ± 0.15 | 91.09 ± 0.17 | 1.59 ± 0.18 | 86.02 ± 0.28 | 87.10 ± 0.30 | 3.34 ± 0.08 | 62.46 ± 1.12 |
| *C4-AD2-ADx1-ADx2 → C7* | 90.41 ± 0.22 | 90.88 ± 0.23 | 1.25 ± 0.17 | 86.15 ± 0.45 | 87.34 ± 0.52 | 4.04 ± 0.58 | 62.55 ± 1.12 |
| *C4-AD2-ADx1-ADx2-ADx4 → C7* | 90.37 ± 0.27 | 90.74 ± 0.27 | 0.99 ± 0.14 | 86.30 ± 0.96 | 87.72 ± 1.09 | 4.66 ± 0.68 | 61.88 ± 1.67 |

Table 9: Performance of JIND-Multi versus JIND and MARS for the *Brain Neurips* scRNA-Seq dataset. Batches column indicate the source batches (left of arrow) and target batch (right of arrow). For JIND, when more than one labeled batch is used, the batches are merged into one.

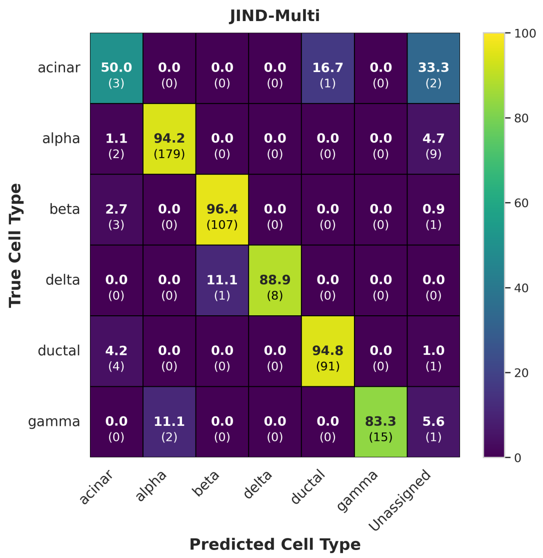

(a)

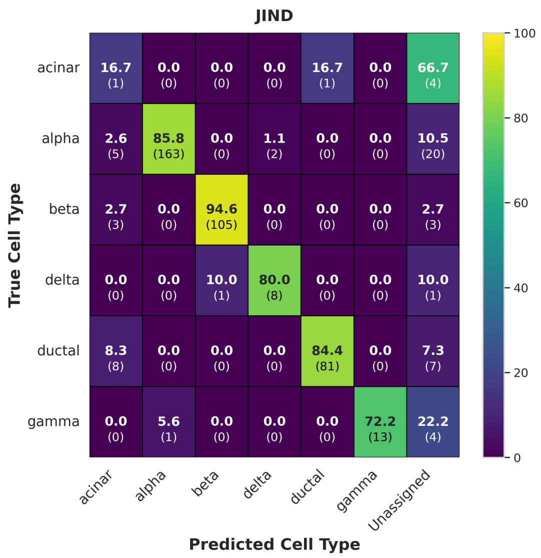

(b)

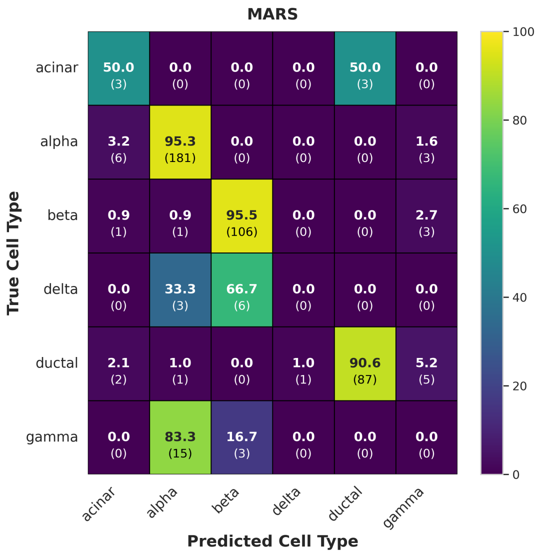

(c)

Figure 8: Confusion matrices showing the prediction accuracies for all cell types in the *Pancreas scRNA-Seq* dataset, using JIND-Multi and MARS trained on *batch 0-1-2*, and JIND trained on *batch 0* with the best trial. JIND-Multi outperforms its predecessor JIND by significantly reducing the number of rejected cells and improving accuracy across all cell types. In comparison to MARS, similar performance is observed for *alpha*, *beta*, and *ductal* cells, while MARS fails to predict the *gamma* and *delta* cell types, which are underrepresented in the source batch.

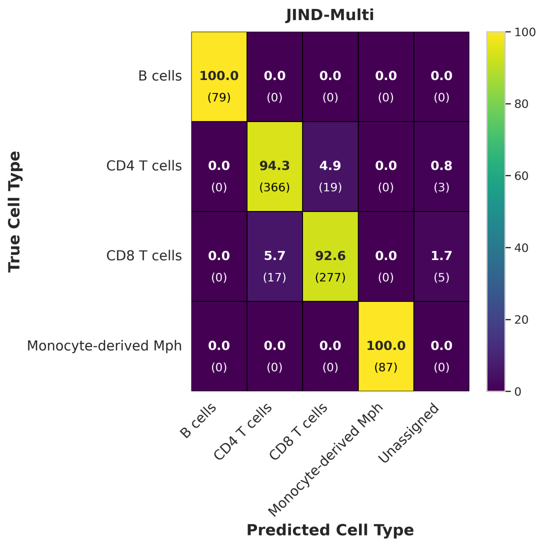

(a)

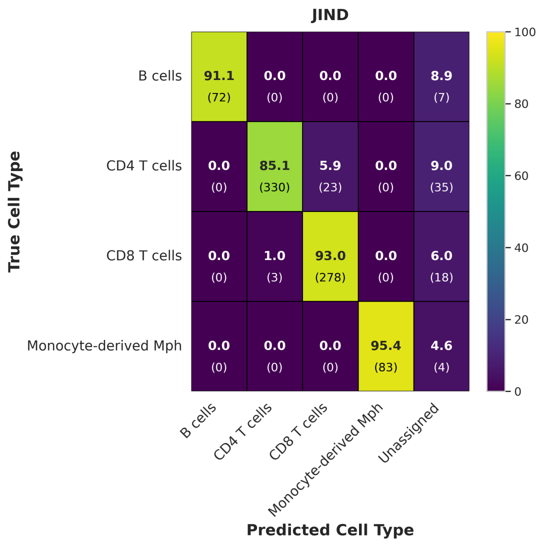

(b)

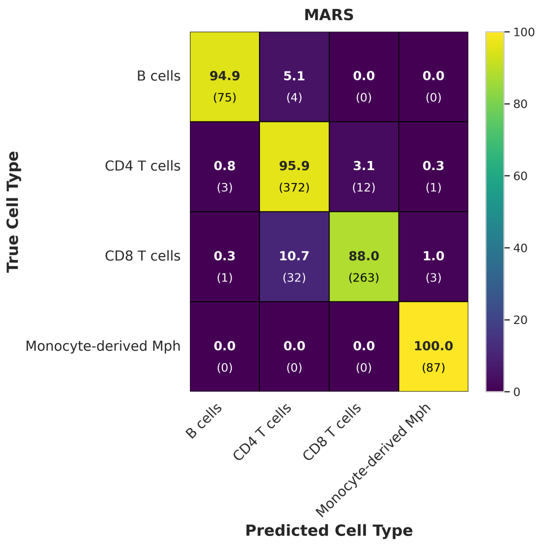

(c)

Figure 9: Confusion matrices showing the prediction accuracies for all cell types on the target batch in *NSCLC Lung scRNA-Seq* dataset. Results include predictions from JIND-Multi and MARS trained on batches *D5-D0-D1-D3-D4-D6*, and JIND trained on batch *D5* with the best trial. JIND-Multi reduces the number of rejected cells compared to JIND, improving accuracy, successfully classifying all B cells and Monocyte-derived Mph. The best run using MARS also improves upon JIND's performance, though there is a higher rate of misclassification among different T cells, as well as between B cells and CD4 T cells.

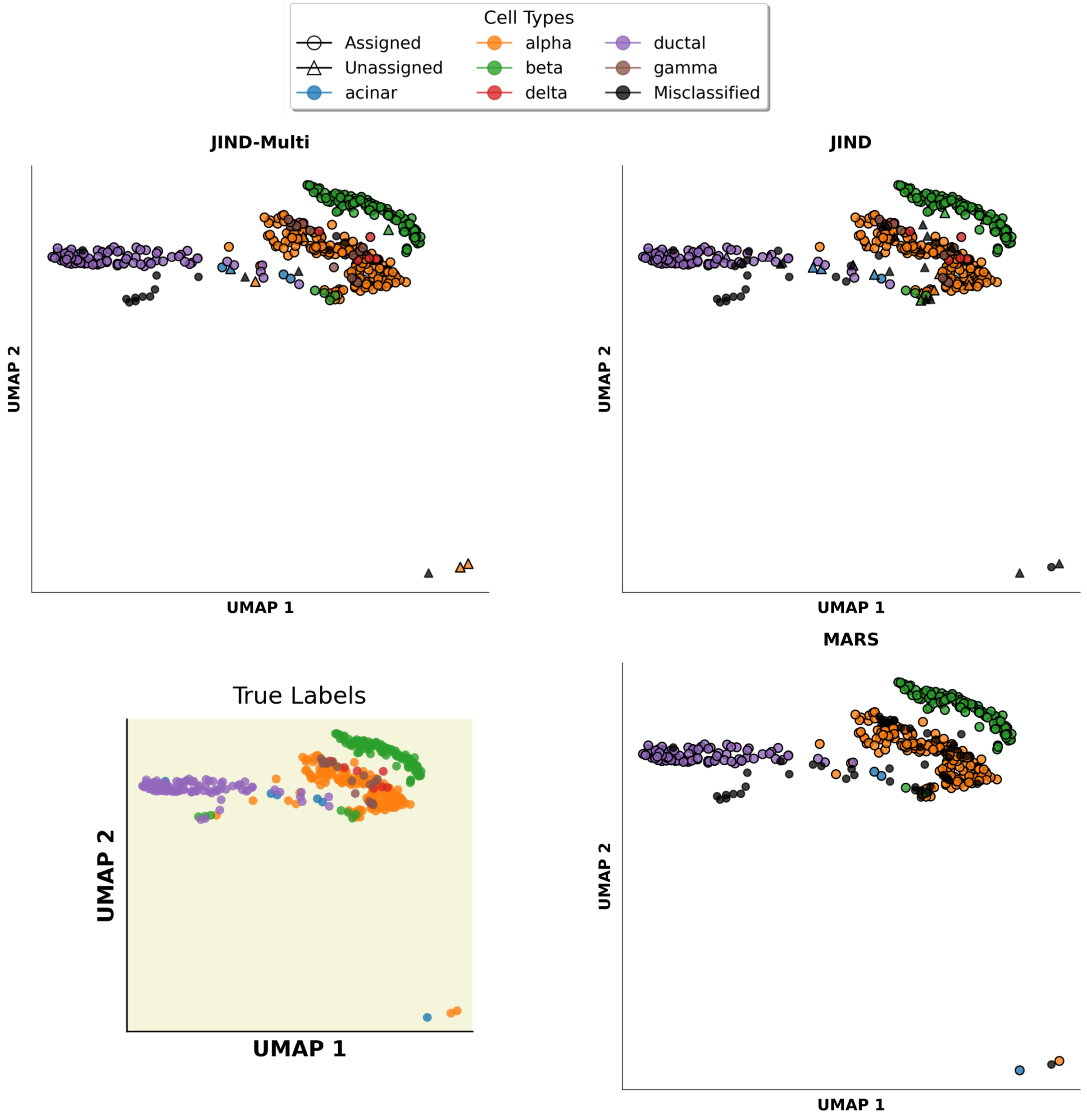

Figure 10: UMAP of the cells' gene expression profiles on the target batch for the *Pancreas scRNA-Seq* dataset with JIND-Multi, JIND and MARS. Cells are colored by cell type and displayed as circles for correct raw predictions from training the model on batches 0-1-2 with the best trial. Misclassified cells are denoted by a black circle, while 'unassigned' cells after filtering with JIND and JIND-Multi are marked with a triangle shape. True label color map is also included. The results illustrate how MARS, unlike JIND-Multi, fails to predict delta and gamma cells. Additionally, out of the 27 incorrect predictions by JIND-Multi, only 13 are classified as errors, as the rest are labeled as unassigned. In contrast, MARS misclassifies 53 cells out of 430.

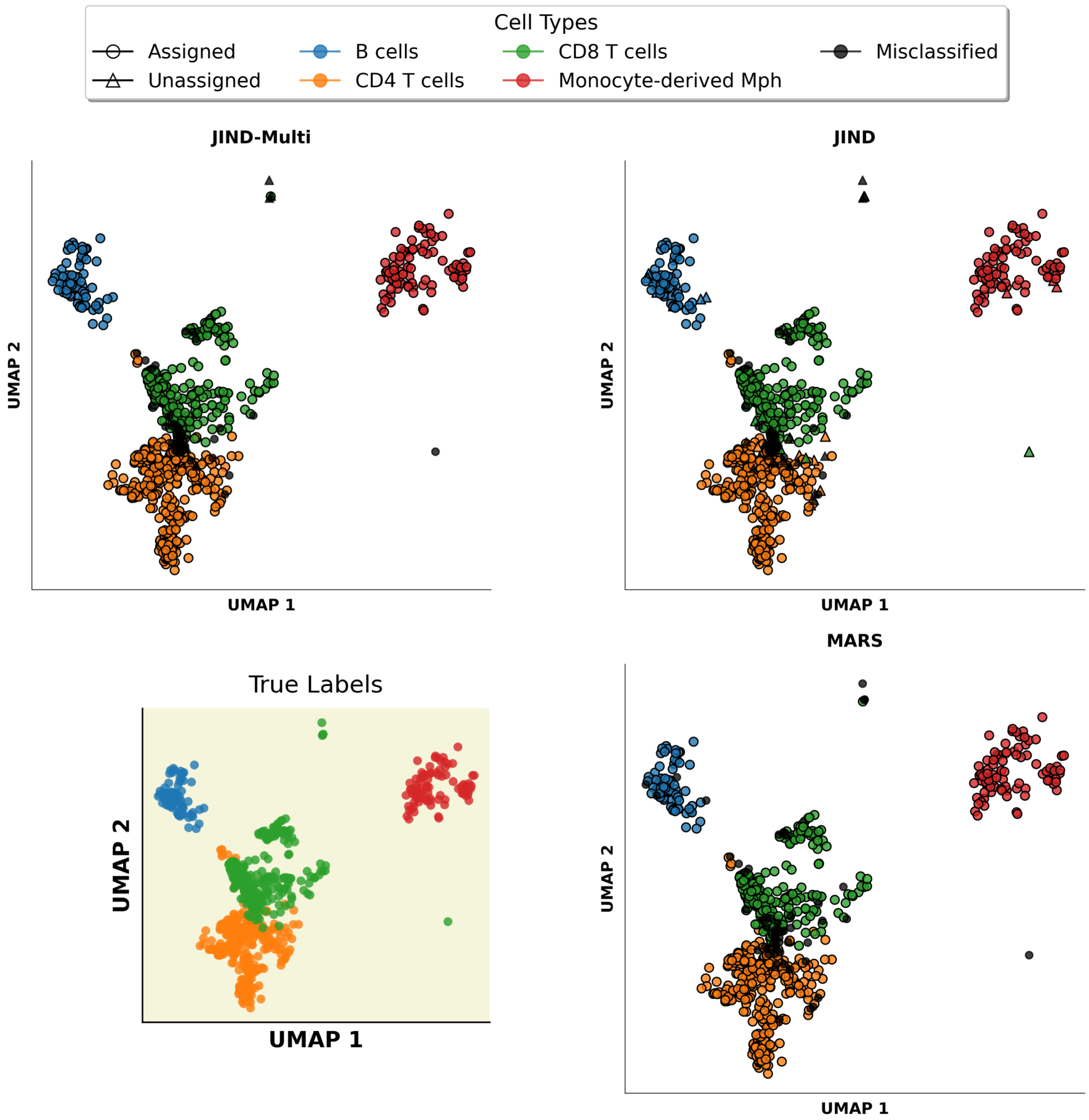

Figure 11: UMAP of the cells' gene expression profiles on the target batch for the *NSCLC Lung scRNA-Seq* dataset with JIND-Multi, JIND and MARS. Cells are colored by cell type and displayed as circles for correct raw predictions from training the model on batches D5-D0-D1-D3-D4-D6 with the best trial. Misclassified cells are denoted by a black circle, while 'unassigned' cells after filtering with JIND and JIND-Multi are marked with a triangle shape. True label color map is also included. The results highlight how monocyte-derived macrophages and B cells, as the most distinct cell types, are perfectly predicted with JIND-Multi, while only in 4 cases do we encounter issues predicting B cells with MARS. Among the various subtypes of T cells, more errors are observed, with MARS specifically misclassifying 52 of these subtypes. In contrast, out of 36 misclassified cells by JIND-Multi, 8 are labeled as unassigned.

##### 3.2 scATAC-Seq experiments

For the annotation of scATAC-Seq data, we compare JIND-Multi with AtacAnnoR [8]. It should be noted that the set-up of AtacAnnoR and JIND-Multi differ substantially, as the former uses annotated scRNA-Seq data as a reference for the annotation of scATAC-Seq data. Additionally, AtacAnnoR does not support the use of several datasets for training nor does it implement cell filtering based on confidence predictions as JIND does. For the comparison, we considered the BMMC scATAC-Seq dataset, as it is the only one for which scRNA-Seq data was also available. We utilized batch s3d3 as target and batch s4d8 as the main source, as in the previous analyses. Both methods were executed 10 times. For AtacAnnoR, we used the scRNA-Seq data from the source batch to infer cell types on the target's scATAC-Seq data. We observe that both methods obtain comparable results.

| Batches | JIND-Multi | | | JIND | | | AtacAnnoR |
|---|---|---|---|---|---|---|---|
| | raw. acc | eff.acc | rej | raw.acc | eff.acc | rej | acc |
| *s4d8 → s3d3* | | | | 89.43 ± 0.00 | 91.04 ± 0.00 | 7.40 ± 0.00 | 89.11 ± 1.23 |
| *s4d8-s1d1 → s3d3* | 88.33 ± 0.41 | 89.41 ± 0.18 | 4.55 ± 0.15 | 90.88 ± 0.00 | 92.27 ± 0.00 | 5.70 ± 0.00 | |
| *s4d8-s1d1-s1d2 → s3d3* | 87.74 ± 0.46 | 88.67 ± 0.47 | 3.92 ± 0.36 | 88.86 ± 0.17 | 89.69 ± 0.30 | 4.52 ± 0.09 | |
| *s4d8-s1d1-s1d2-s1d3 → s3d3* | 87.21 ± 0.98 | 87.89 ± 0.95 | 3.20 ± 0.53 | 89.88 ± 0.77 | 91.06 ± 0.70 | 4.23 ± 0.86 | |
| *s4d8-s1d1-s1d2-s1d3-s2d1 → s3d3* | 87.13 ± 0.74 | 87.62 ± 0.73 | 2.25 ± 0.24 | 89.95 ± 0.71 | 90.66 ± 0.57 | 2.63 ± 0.85 | |
| *s4d8-s1d1-s1d2-s1d3-s2d1-s2d4 → D2* | 87.82 ± 0.54 | 88.28 ± 0.58 | 1.66 ± 0.22 | 90.47 ± 0.60 | 91.08 ± 0.58 | 1.97 ± 0.52 | |
| *s4d8-s1d1-s1d2-s1d3-s2d1-s2d4-s2d5 → D2* | 87.56 ± 0.60 | 87.83 ± 0.59 | 1.36 ± 0.51 | 90.59 ± 0.43 | 91.14 ± 0.40 | 1.66 ± 0.45 | |
| *s4d8-s1d1-s1d2-s1d3-s2d1-s2d4-s2d5-s3d10 → D2* | 87.54 ± 0.99 | 87.82 ± 0.99 | 1.15 ± 0.25 | 90.85 ± 0.52 | 91.30 ± 0.43 | 1.71 ± 0.48 | |
| *s4d8-s1d1-s1d2-s1d3-s2d1-s2d4-s2d5-s3d10-s4d1 → D2* | 87.71 ± 0.83 | 87.91 ± 0.83 | 0.81 ± 0.18 | 90.73 ± 0.70 | 91.24 ± 0.69 | 1.59 ± 0.28 | |

Table 10: Performance of JIND, JIND-Multi, and AtacAnnoR on the *BMMC* scATAC-Seq dataset. Batches column indicate the source batches (left of arrow) and target batch (right of arrow). For JIND, when more than one labeled batch is used, the batches are merged into one. AtacAnnoR's accuracy is shown in the last column.

| Batches | JIND-Multi | | | JIND | | |
|---|---|---|---|---|---|---|
| | raw. acc | eff.acc | rej | raw.acc | eff.acc | rej |
| *heart sample 39 → heart sample 14* | | | | 77.21 ± 0.00 | 78.04 ± 0.00 | 2.20 ± 0.00 |
| *heart sample 39-heart sample 32 → heart sample 14* | 77.45 ± 0.08 | 77.72 ± 0.04 | 0.76 ± 0.06 | 77.01 ± 0.00 | 77.53 ± 0.00 | 1.60 ± 0.00 |

Table 11: Performance of JIND and JIND-Multi on the *Fetal Heart* scATAC-Seq dataset. Batches column indicate the source batches (left of arrow) and target batch (right of arrow). For JIND, when more than one labeled batch is used, the batches are merged into one.

| Batches | JIND-Multi | | | JIND | | |
|---|---|---|---|---|---|---|
| | raw. acc | eff.acc | rej | raw.acc | eff.acc | rej |
| *kidney s3 → kidney s67* | | | | 67.81 ± 0.00 | 68.98 ± 0.00 | 5.70 ± 0.00 |
| *kidney s3-kidney s34 → kidney s67* | 69.12 ± 0.36 | 69.46 ± 0.47 | 1.11 ± 0.38 | 70.03 ± 0.00 | 70.93 ± 0.00 | 4.10 ± 0.00 |
| *kidney s3-kidney s34-kidney s65 → kidney s67* | 69.03 ± 1.06 | 69.32 ± 1.26 | 1.31 ± 0.36 | 70.67 ± 0.29 | 71.92 ± 0.41 | 4.16 ± 0.09 |

Table 12: Performance of JIND and JIND-Multi on the *Fetal Kidney* scATAC-Seq dataset. Batches column indicate the source batches (left of arrow) and target batch (right of arrow). For JIND, when more than one labeled batch is used, the batches are merged into one.

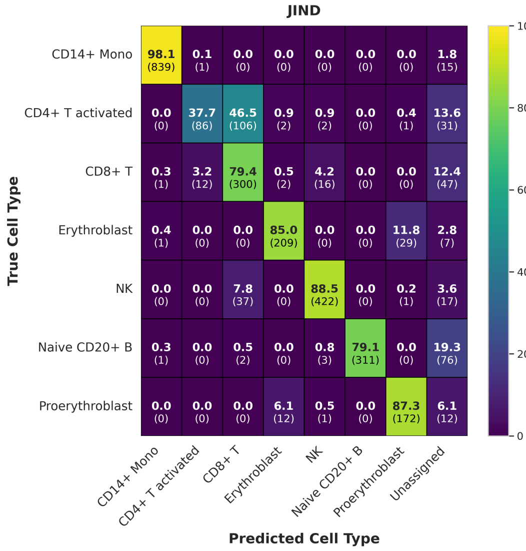

(a)

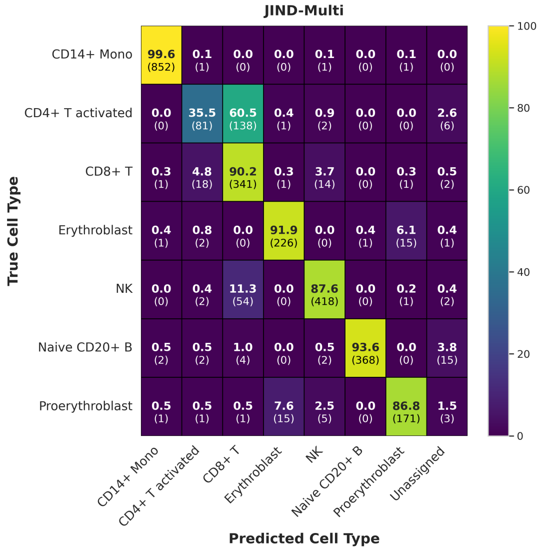

(b)

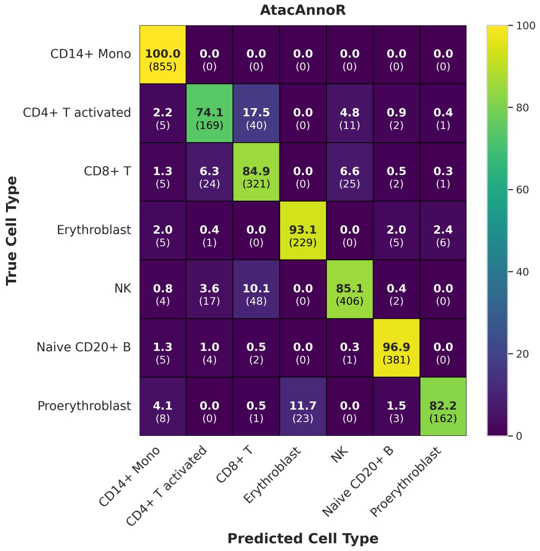

(c)

Figure 12: Confusion matrices showing the prediction accuracies for all cell types in the *BMMC scATAC-Seq* dataset, with results from JIND-Multi trained on batches *s4d8-s1d1-s1d2-s1d3-s2d1-s2d4-s2d5-s3d10-s4d1* and AtacAnnoR and JIND trained on batch *s4d8* with the best trial. JIND-Multi drastically reduces the percentage of rejected cells compared to JIND and significantly improves classification, especially for CD4+ T activated, CD8+ T, erythroblast, and Naive CD20+ B cells, achieving around 90% accuracy for these cell types. AtacAnnoR, with a performance similar to JIND-Multi, is able to better distinguish between CD4+ T activated and CD8+ T cells.

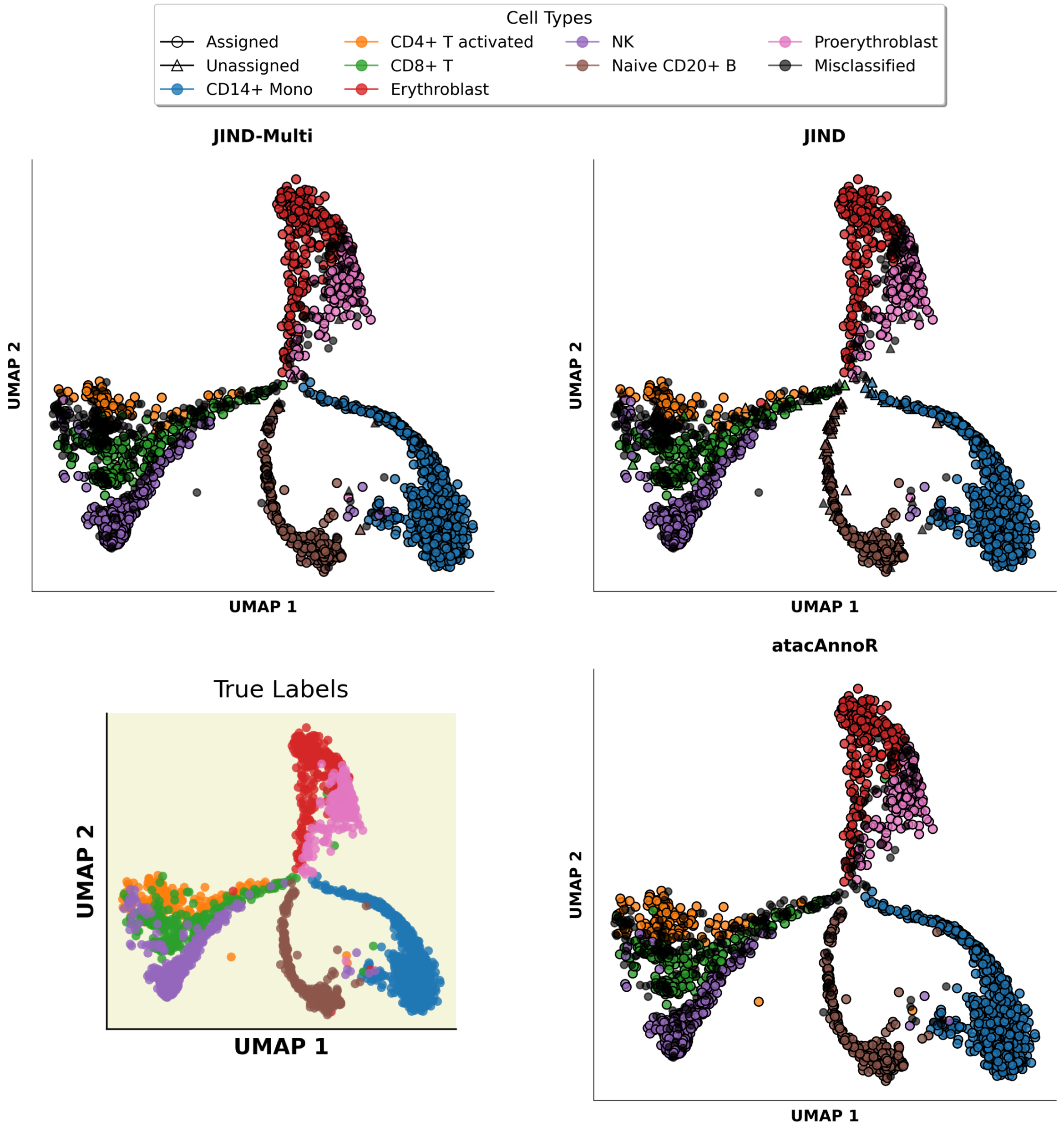

Figure 13: UMAP of the cells' gene expression profiles on the target batch for the *BMMC scATAC-Seq* dataset with JIND-Multi, JIND and atacAnnoR. Cells are colored by cell type and displayed as circles for correct raw predictions from training the model on batches *s4d8-s1d1-s1d2-s1d3-s2d1-s2d4-s2d5-s3d10-s4d1* with JIND-Multi and AtacAnnoR and JIND trained on batch s4d8 with the best trial. Misclassified cells are denoted by a black circle, while 'unassigned' cells after filtering with JIND and JIND-Multi are marked with a triangle shape. True label color map is also included.

#### 3.3 Processing speed

| Batches | JIND-Multi time (s) | JIND time (s) |
|---|---|---|
| *0 → 3* | | *61* |
| *0-1 → 3* | *156* | *54* |
| *0-1-2 → 3* | *266* | *59* |

Table 13: Processing speed of JIND and JIND-Multi on the *Pancreas* scRNA-Seq dataset. Batches column indicate the source batches (left of arrow) and target batch (right of arrow). For JIND, when more than one labeled batch is used, the batches are merged into one.

| Batches | JIND-Multi time (s) | JIND time (s) |
|---|---|---|
| *D5 → D2* | | *50* |
| *D5-D0 → D2* | *133* | *56* |
| *D5-D0-D1 → D2* | *266* | *178* |
| *D5-D0-D1-D3 → D2* | *394* | *64* |
| *D5-D0-D1-D3-D4 → D2* | *508* | *71* |
| *D5-D0-D1-D3-D4-D6 → D2* | *756* | *140* |

Table 14: Processing speed of JIND and JIND-Multi on the *NSCLC Lung* scRNA-Seq dataset. Batches column indicate the source batches (left of arrow) and target batch (right of arrow). For JIND, when more than one labeled batch is used, the batches are merged into one.

| Batches | JIND-Multi time (s) | JIND time (s) |
|---|---|---|
| *C4 → C7* | | *403* |
| *C4-AD2 → C7* | *841* | *352* |
| *C4-AD2-ADx1 → C7* | *1370* | *358* |
| *C4-AD2-ADx1-ADx2 → C7* | *1975* | *399* |
| *C4-AD2-ADx1-ADx2-ADx4 → C7* | *2717* | *434* |

Table 15: Processing speed of JIND and JIND-Multi on the *Brain Neurips* scRNA-Seq dataset. Batches column indicate the source batches (left of arrow) and target batch (right of arrow). For JIND, when more than one labeled batch is used, the batches are merged into one.

| Batches | JIND-Multi time (s) | JIND time (s) | AtacAnnoR time (s) |
|---|---|---|---|
| *s4d8 → s3d3* | | *84* | *44* |
| *s4d8-s1d1 → s3d3* | *169* | *76* | |
| *s4d8-s1d1-s1d2 → s3d3* | *271* | *88* | |
| *s4d8-s1d1-s1d2-s1d3 → s3d3* | *401* | *86* | |
| *s4d8-s1d1-s1d2-s1d3-s2d1 → s3d3* | *557* | *105* | |
| *s4d8-s1d1-s1d2-s1d3-s2d1-s2d4 → s3d3* | *775* | *102* | |
| *s4d8-s1d1-s1d2-s1d3-s2d1-s2d4-s2d5 → s3d3* | *1072* | *111* | |
| *s4d8-s1d1-s1d2-s1d3-s2d1-s2d4-s2d5-s3d10 → s3d3* | *1485* | *122* | |
| *s4d8-s1d1-s1d2-s1d3-s2d1-s2d4-s2d5-s3d10-s4d1 → s3d3* | *1995* | *140* | |

Table 16: Processing speed of JIND, JIND-Multi, and AtacAnnoR on the *BMMC* scATAC-Seq dataset. Batches column indicates the source batches (left of arrow) and target batch (right of arrow). For JIND, when more than one labeled batch is used, the batches are merged into one.

| Batches | JIND-Multi time (s) | JIND time (s) |
|---|---|---|
| *heart sample 39 → heart sample 14* | | *141* |
| *heart sample 39-heart sample 32 → heart sample 14* | *367* | *113* |

Table 17: Processing speed of JIND and JIND-Multi on the *Fetal Heart* scATAC-Seq dataset. Batches column indicate the source batches (left of arrow) and target batch (right of arrow). For JIND, when more than one labeled batch is used, the batches are merged into one.

| Batches | JIND-Multi time (s) | JIND time (s) |
|---|---|---|
| *kidney s3 → kidney s67* | | *59* |
| *kidney s3-kidney s34 → kidney s67* | *176* | *67* |
| *kidney s3-kidney s34-kidney s6 → kidney s67* | *291* | *72* |

Table 18: Processing speed of JIND and JIND-Multi on the *Fetal Kidney* scATAC-Seq dataset. Batches column indicate the source batches (left of arrow) and target batch (right of arrow). For JIND, when more than one labeled batch is used, the batches are merged into one.

#### 3.4 Differential Expression Analysis

For each scRNA-Seq dataset and the *BMMC scATAC-Seq* dataset, we performed a differential expression analysis of genes using *scanpy* [8]. This analysis involved calculating a Wilcoxon test for each cluster (cell type) against the remaining cells to identify differentially expressed genes. We utilized the AnnData object, with the *cells x genes* transformed data, which included only the selected batches relevant to our study (see Figure S14-15-16).

.

##### 3.4.1 Compare specific clusters

In addition, we conducted another differential expression (DE) analysis using the *t-test_overestim_var* method. We focused on similar cell-types that posed challenges with the MARS and JIND-Multi-JIND analyses. For each combination of cell types, one type was selected for analysis while the other served as a reference (see Figure S19-22)

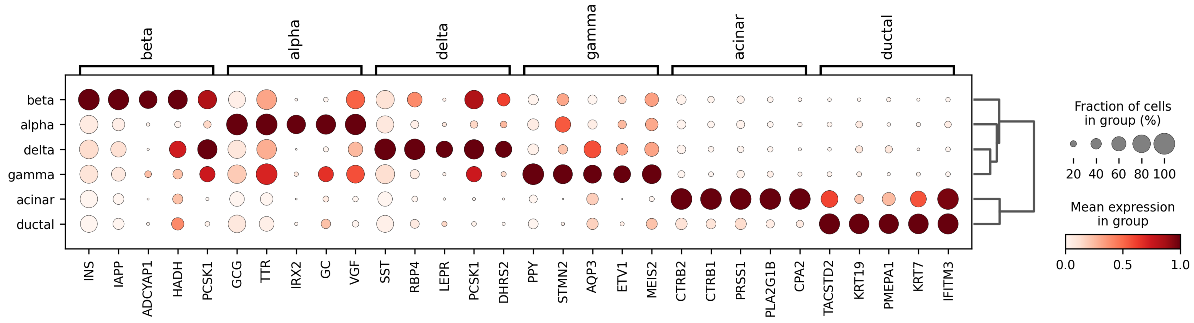

Figure 14: Top 5 differentially expressed genes in *Pancreas scRNA-Seq* dataset for each cell-type. The color intensity indicates the mean expression percentage, while the size of the circles represents the fraction of cells expressing each gene.

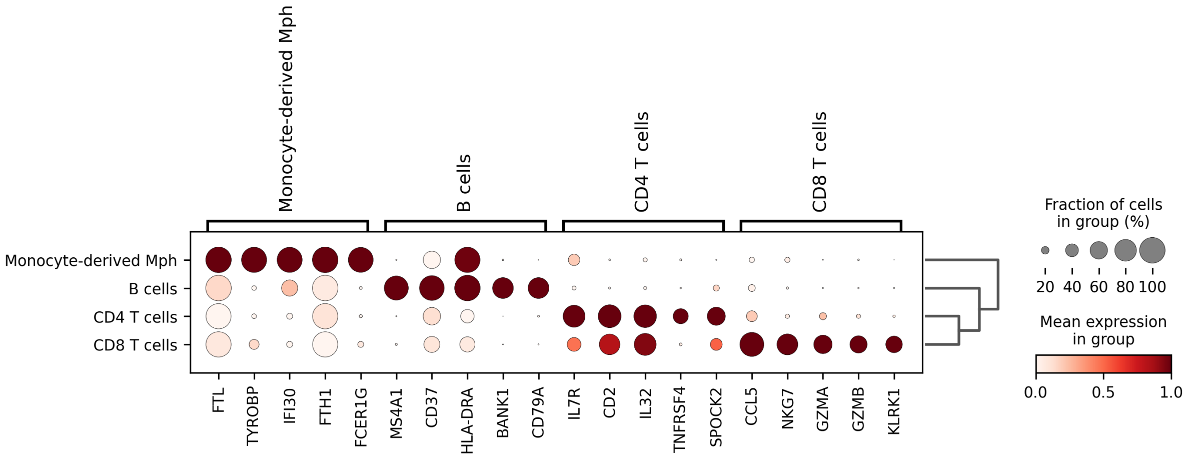

Figure 15: Top 5 differentially expressed genes in *Lung scRNA-Seq* dataset for each cell-type. The color intensity indicates the mean expression percentage, while the size of the circles represents the fraction of cells expressing each gene.

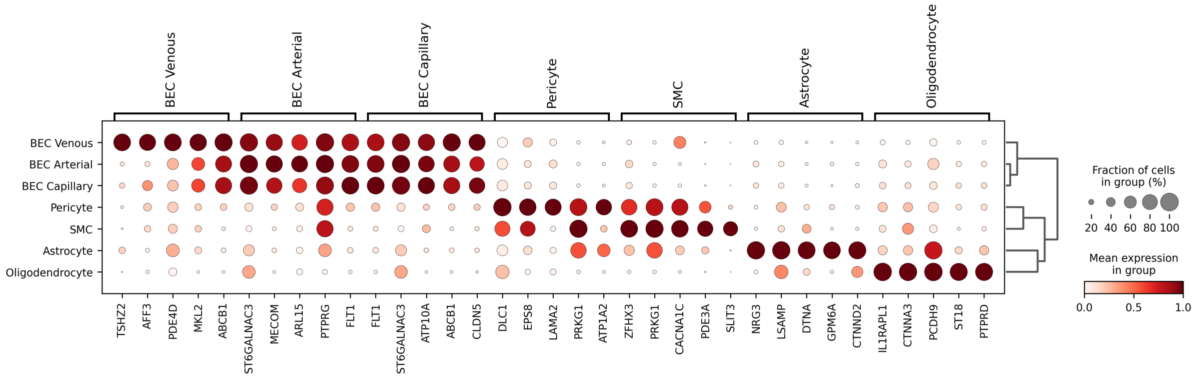

Figure 16: Top 5 differentially expressed genes in *Brain Neurips scRNA-Seq* dataset for each cell-type. The color intensity indicates the mean expression percentage, while the size of the circles represents the fraction of cells expressing each gene.

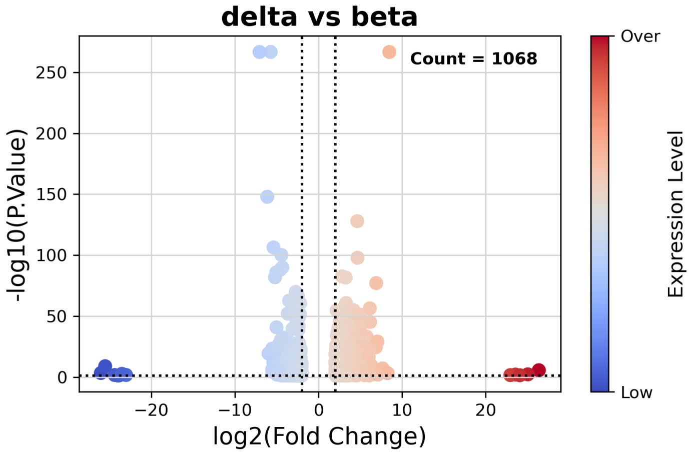

Figure 17: Volcano plot comparing gene expression levels between delta and beta groups in *Pancreas scRNA-Seq* dataset . The x-axis represents the *log2(Fold Change)* of expression, while the y-axis displays the $-\log_{10}(P\text{-}Value)$. Only genes that meet the defined significance thresholds are plotted, with dotted lines indicating thresholds for fold change ($|\log_2(\text{FC})| > 2$) and adjusted p-value ($P < 0.05$). A total of 1,068 differentially expressed genes are identified, color-coded to represent expression levels: blue for ”Low” expression and red for ”Over” expression.

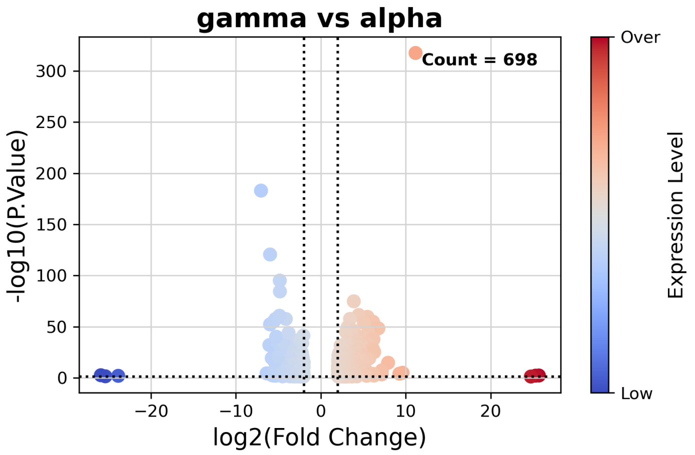

Figure 18: Volcano plot comparing gene expression levels between gamma and alpha groups in *Pancreas scRNA-Seq* dataset . The x-axis represents the *log2(Fold Change)* of expression, while the y-axis displays the $-\log_{10}(P\text{-}Value)$. Only genes that meet the defined significance thresholds are plotted, with dotted lines indicating thresholds for fold change ($|\log_2(\text{FC})| > 2$) and adjusted p-value ($P < 0.05$). A total of 698 differentially expressed genes are identified, color-coded to represent expression levels: blue for ”Low” expression and red for ”Over” expression.

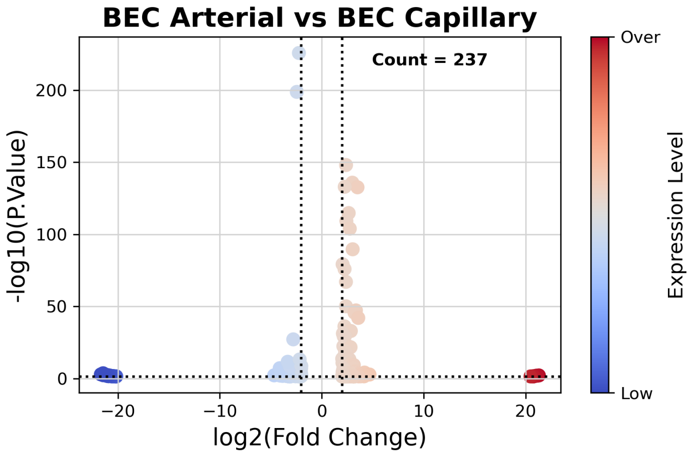

(a)

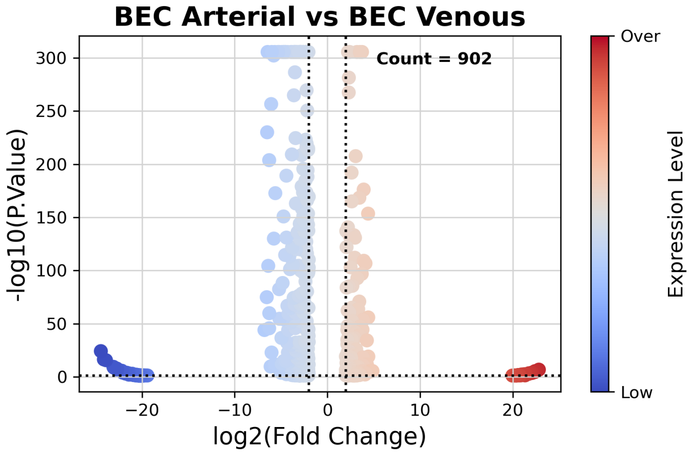

(b)

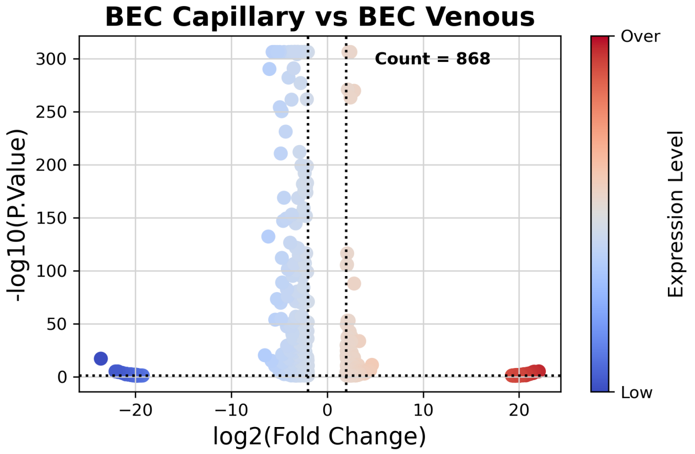

(c)

Figure 19: Volcano plots comparing gene expression levels in *Brain Neurips scRNA-Seq* dataset across three groups: BEC Arterial vs. BEC Capillary (a), BEC Arterial vs. BEC Venous (b), and BEC Capillary vs. BEC Venous (c). Differential expression analysis was performed using the *t-test_overestim_var* method. The x-axis represents the *log2(Fold Change)* of expression, while the y-axis displays the $-\log_{10}(P\text{-}Value)$. Only genes that meet the defined significance thresholds are plotted. Dotted lines indicate significance thresholds for fold change ($|\log_2(\text{FC})| > 2$) and adjusted p-value ($P < 0.05$). Differentially expressed gene are color-coded according to their expression levels, with blue indicating "Low" expression and red indicating "Over" expression. The total number of differentially expressed genes is annotated in each plot. Notably, BEC Arterial and BEC Capillary exhibit high similarity, with only 237 differentially expressed genes.

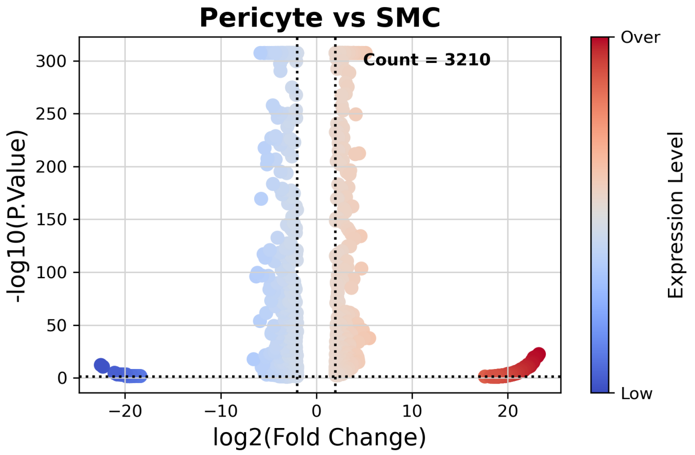

Figure 20: Volcano plot comparing gene expression levels in *Brain Neurips scRNA-Seq* dataset between Pericyte cells and SMC cells, focusing only on genes that meet the defined significance thresholds. The x-axis represents the *log2(Fold Change)* of expression, while the y-axis displays the $-\log_{10}(P\text{-}Value)$. Dotted lines indicate significance thresholds for fold change ($|\log_2(\text{FC})| > 2$) and adjusted p-value ($P < 0.05$). A total of 3210 differentially expressed genes are identified, with color coding to represent expression levels: blue for "Low" expression and red for "Over" expression.

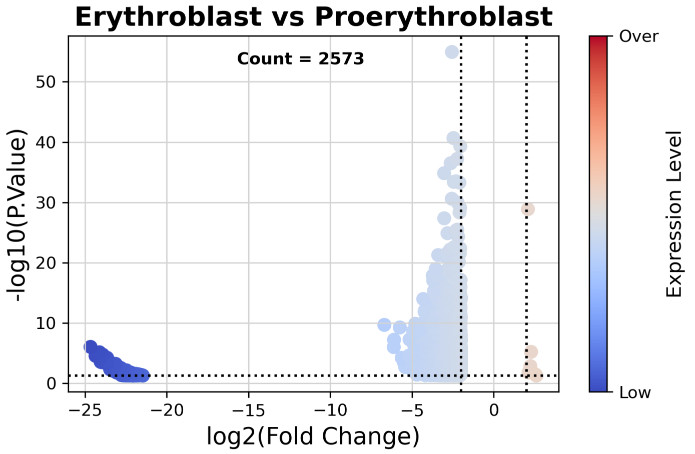

Figure 21: Volcano plot comparing gene expression levels in *BMMC scATAC-Seq* dataset between Erythroblast cells and Proerythroblast cells, focusing only on genes that meet the defined significance thresholds. The x-axis represents the *log2(Fold Change)* of expression, while the y-axis displays the $-\log_{10}(P\text{-}Value)$. Dotted lines indicate significance thresholds for fold change ($|\log_2(\text{FC})| > 2$) and adjusted p-value ($P < 0.05$). A total of 2,573 differentially expressed genes are identified, with color coding to represent expression levels: blue for "Low" expression and red for "Over" expression.

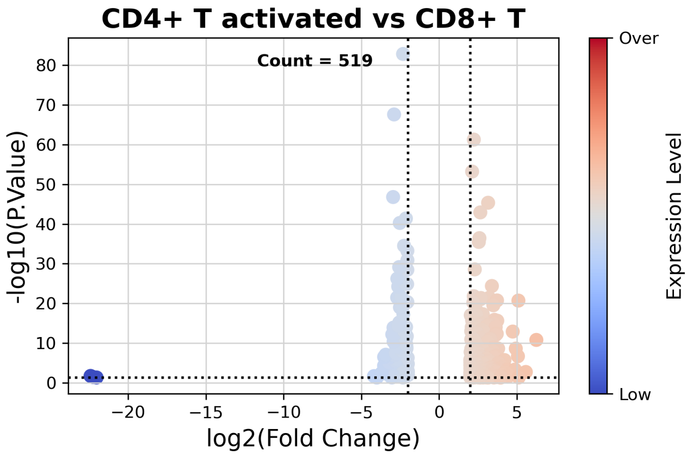

Figure 22: Volcano plot comparing gene expression levels in *BMMC scATAC-Seq* dataset between CD4+ activated T cells and CD8+ T cells, focusing only on genes that meet the defined significance thresholds. The x-axis represents the *log2(Fold Change)* of expression, while the y-axis displays the $-\log_{10}(P\text{-}Value)$. Dotted lines indicate significance thresholds for fold change ($|\log_2(\mathrm{FC})| > 2$) and adjusted p-value ($P < 0.05$). A total of 519 differentially expressed genes are identified, with color coding to represent expression levels: blue for "Low" expression and red for "Over" expression.

[4] Kaiming He, Xiangyu Zhang, Shaoqing Ren, and Jian Sun. Deep residual learning for image recognition. *CoRR*, abs/1512.03385, 2015.

[5] Leland McInnes, John Healy, Nathaniel Saul, and Lukas Großberger. Umap: Uniform manifold approximation and projection. *Journal of Open Source Software*, 3(29):861, 2018.

[6] Mauro J Muraro, Gitanjali Dharmadhikari, Dominic Grün, Niels Groen, Thomas Dielen, Erik Jansen, Leon van Gurp, Marten A Engelse, Françoise Carlotti, Eelco J de Koning, and Alexander van Oudenaarden. A single-cell transcriptome atlas of the human pancreas. *Cell Systems*, 3(4):385–394.e3, 2016.

[7] Åsa Segerstolpe, Athanasia Palasantza, Pernilla Eliasson, Eva M Andersson, Anna C Andréasson, Xiaoyan Sun, Simone Picelli, Alan Sabirsh, Mattias Clausen, Magnus K Bjursell, Donald M Smith, Michael Kasper, Carl Ämmälä, and Rickard Sandberg. Single-cell transcriptome profiling of human pancreatic islets in health and type 2 diabetes. *Cell Metabolism*, 24(4):593–607, 2016.

[8] Florian A Wolf, Philipp Angerer, and Fabian J Theis. Scanpy: large-scale single-cell gene expression data analysis. *Genome Biology*, 19(1):15, 2018.
